# GIS-FA: An approach to integrate thematic maps, factor-analytic and envirotyping for cultivar targeting

**DOI:** 10.1101/2023.07.15.549137

**Authors:** Maurício S. Araújo, Saulo F. S. Chaves, Luiz A. S. Dias, Filipe M. Ferreira, Guilherme R. Pereira, André R. G. Bezerra, Rodrigo S. Alves, Alexandre B. Heinemann, Flávio Breseghello, Pedro C. S. Carneiro, Matheus D. Krause, Germano Costa-Neto, Kaio O. G. Dias

**Author notes:** These authors contributed equally to this work (Maurício S. Araújo, Saulo F. S. Chaves).

## Abstract

**Key message: We propose an enviromics prediction model for cultivar recommendation based on thematic maps for decision-makers**.

Parsimonious methods that capture genotype-by-environment interaction (GEI) in multi-environment trials (MET) are important in breeding programs. Understanding the causes and factors of GEI allows the utilization of genotype adaptations in the target population of environments through environmental features and Factor-Analytic (FA) models. Here, we present a novel predictive breeding approach called GIS-FA that integrates geographic information systems (GIS) techniques, FA models, Partial Least Squares (PLS) regression, and Enviromics to predict phenotypic performance in untested environments. The GIS-FA approach allows: (i) predict the phenotypic performance of tested genotypes in untested environments; (ii) select the best-ranking genotypes based on their overall performance and stability using the FA selection tools; (iii) draw thematic maps showing overall or pairwise performance and stability for decision-making. We exemplify the usage of GIS-FA approach using two datasets of rice [*Oryza sativa* (L.)] and soybean [*Glycine max* (L.) Merr.] in MET spread over tropical areas. In summary, our novel predictive method allows the identification of new breeding scenarios by pinpointing groups of environments where genotypes have superior predicted performance and facilitates/optimizes the cultivar recommendation by utilizing thematic maps.

## 1 Introduction

Crossover interaction refers to changes in the ranking of genotypes caused by the lack of genotypic correlation between environments, which is the most critical source of genotype-by-environment interaction (GEI) for plant breeders (Cooper and Delacy, 1994; Crossa et al, 2004). Cultivar development programs of staple crops evaluate experimental genotypes (i.e., prior to release) in multi-environmental trials (MET) to (*i*) depict GEI patterns for future cultivar placement and (*ii*) increase the accuracy of selection. Therefore, analytical methods that fully explore the GEI patterns from MET are needed for decision-making (Malosetti et al, 2013; van Eeuwijk et al, 2016; Dias et al, 2022; Tolhurst et al, 2022).

The first attempt to consider the GEI in plant breeding was proposed by Yates and Cochran (1938), who decomposed the part due to the interaction from the total phenotypic variation. Later, Finlay and Wilkinson (1963) used marginal environmental means as independent variables in the regression analysis to depict GEI, and several approaches were developed in that framework (Eberhart and Russell, 1966; Li et al, 2018). Multivariate techniques such as the additive main effects and multiplicative interaction (AMMI) (Gauch JR and Zobel, 1997) and the genotype plus GEI (GGE) biplot (Yan et al, 2000) have also been extensively used (Yan et al, 2007; Balestre et al, 2009; Silva et al, 2021). Further model expansions were made possible by the development of the linear mixed model equations (Henderson, 1949, 1950), in which the covariance between relatives and environments could be incorporated, and assumptions such as homogeneous residual variances could be relaxed (Piepho et al, 2008). Factor-analytic (FA) mixed models (Piepho, 1997; Smith et al, 2001) can be employed to explore the covariance between environments. These models offer the flexibility to account for heterogeneous genotypic (or genetic) covariances between environments using a few latent variables known as factors (*K*). In addition to the overall (i.e., across environments) and conditional (i.e., within environments) performance, metrics such as stability and sensitivity can also be computed from FA models to facilitate the decision-making process (Stefanova and Buirchell, 2010; Cullis et al, 2014; Dias et al, 2018; Smith and Cullis, 2018; Smith et al, 2021).

An extension to statistical models that deal with GEI is the incorporation of environmental information, such as physical and chemical soil properties, as well as environmental features like temperature and rainfall precipitation (Bakare et al, 2022). The advantages of incorporating environmental features in a prediction model are (*i*) the ability to untangle environmental determinants and the crossover GEI main drivers, and (*ii*) to predict phenotypic performance in yet-to-be-seen environments (Sae-Lim et al, 2014; Oliveira et al, 2020; Tolhurst et al, 2022). Furthermore, the delineation of homogeneous groups of environments based on their environmental similarity facilitates resource optimization and the identification of mega-environments (Wood, 1976; Denis, 1988; Van Eeuwijk and Elgersma, 1993; Millet et al, 2019; Costa-Neto et al, 2021c; Krause et al, 2022). Therefore, advances in computational resources along with the development of geographic information systems (GIS) techniques are essential to design novel prediction strategies in MET (Cooper and Messina, 2021; Rogers et al, 2021; Cooper et al, 2022; Diepenbrock et al, 2022).

GIS techniques have been defined as computer-based systems used for analyzing and interpreting spatially referenced information (Coppock and Rhind, 1991), and are powerful tools in the integration of genetics and environmental information (Chapman et al, 1996; Beebe et al, 1997; Guarino et al, 2002; Jarquíin et al, 2014; Hernández et al, 2019; Costa-Neto and Fritsche-Neto, 2021). For example, Annicchiarico et al (2006) identified repeatable genotype-by-location interactions using GIS-based models for cultivar recommendation for durum wheat in Algeria. Costa-Neto et al (2020) applied a GIS-based tool with factorial regression to model spatial trends and draw thematic maps of yield performance to upland rice in Brazil; and Costa-Neto et al (2021b) integrated GIS techniques with nonlinear kernels to model additive, dominance, and GEI effects. All the mentioned techniques follow under the umbrella of “envirotypic-assisted selection”, which integrates genomic with environmental data to improve the accuracy of selection in plant breeding programs (Resende et al, 2021).

The combination of statistics, quantitative genetics, and GIS techniques enabled the introduction of the field of Enviromics in the plant breeding community (Cooper et al, 2014; Xu, 2016; Costa-Neto and Fritsche-Neto, 2021). Coupled with knowledge from plant ecophysiology, it aims to describe how the environment impact plant development and phenotypic plasticity of important agronomic traits (Costa-Neto and Fritsche-Neto, 2021). Accordingly, envirotypes are all sources of environmental variations related to plant development that can act as environmental markers in statistical genetics models to predict genotypic effects in non-evaluated environments (Xu, 2016; Resende et al, 2021). However, the integration of phenotypic and genomic data with environmental features can generate two statistical problems: predictors (*p*) might have a high correlation resulting in multicollinearity, and the curse of dimensionality when the number of observations (*n*) is smaller than *p* (*n* ≪ *p*). In these situations, methods such as partial least squares (PLS) that combine features from principal components analysis and multiple regression (Wold et al, 2001) and Bayesian factor analytic model (Nuvunga et al, 2019) can be applied to identify linear combinations of *p* that capture the underlying structure of the data (Montesinos-López et al, 2022a,b).

Here, we present a novel predictive breeding approach called GIS-FA that combines FA, PLS, and enviromics to predict the phenotypic performance of experimental genotypes in untested environments. The GIS-FA uses environmental information collected from GIS tools to predict the factor loadings of untested environments via PLS, where the estimated factor loadings from the observed environments are used as training set. The empirical best linear unbiased predicted values (eBLUPs) of genotypic means in untested environments are then calculated as the linear combination of the predicted loadings via PLS and genotypic scores from the FA models. We hypothesize the GIS-FA model has a higher prediction accuracy when compared to a PLS model being trained with eBLUPs within observed environments (henceforth called GIS-GGE). We tested this hypothesis in two MET datasets from Brazil: rice trials located in the Brazilian Savanna (Cerrado) and in the Amazon rainforest, and soybean trials located in the state of Mato Grosso do Sul. Thus, this study intends to (*i*) propose the GIS-FA methodology for predicting genotypes’ performance in untested environments, and compare its predictive ability with GIS-GGE methodology; (*ii*) apply GIS-FA to select the best-ranking genotypes based on their overall performance and stability using the FA selection tools; and (*iii*) draw thematic maps that illustrate the genotype’s performance across environments in the breeding zone.

## 2 Material and Methods

### 2.1 Phenotypic data

We exemplify the GIS-FA model using two datasets from MET spread over tropical areas in Brazil. These trials have been used for decisions on cultivar release by public and proprietary breeding organizations. The soybean dataset contains three years of field trials conducted in the state of Mato Grosso do Sul (triangles in Figure 1), whereas the rice dataset has two years of field trials spread over eight states (circles in Figure 1). Note how the elevation varies across the studied area (Figure 1a). This factor, along with the latitude and longitude, dictates the changes in both weather and soil conditions, represented by the Köppen-Geiger classification (Alvares et al, 2013) in Figure 1b, and the Brazilian Soil Classification System (Santos, 2018) in Figure 1c. In both datasets, there were field trials planted in the same location and year, but with different planting seasons. Thus, henceforth, the term “environment” refers to the combination of location, year, and planting season. Another common characteristic between both datasets is the fact that not all genotypes were evaluated in all environments (Supplementary Figure 1). This has three main reasons: i) seed availability; ii) low-performing lines were discarded at the end of each agricultural year; and iii) cultivars/genotypes from partner breeding programs were included for evaluation in the TPE.

**Fig. 1.**
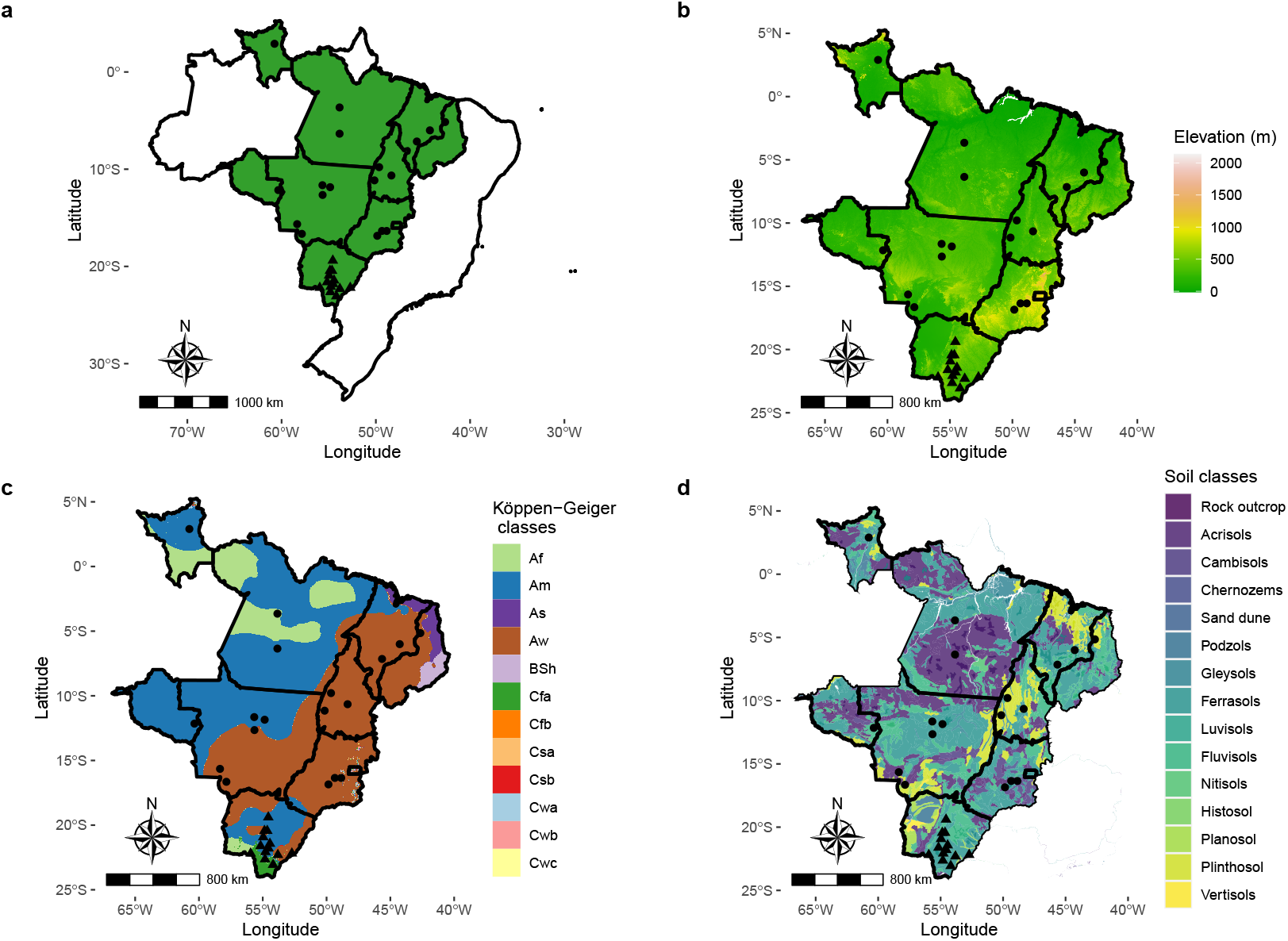
Maps of the studied area: a) Map of Brazil with rice or soy trials b) elevation in meters, c) Köppen-Geiger classification (Alvares et al, 2013), and d) Brazilian soil classification adapted to FAO classification (Santos, 2018; Fao, 2014). Triangles and points within the maps represent the geographic locations of the soybean and rice trials, respectively. In the lower right corner, the states where these trials were conducted are highlighted.

#### 2.1.1 Rice dataset

The rice dataset is composed of 80 purelines developed by the Brazilian Agricultural Research Corporation (Embrapa Rice and Beans). These purelines plus three commercial cultivars were evaluated for their value of cultivation and use (VCU) in 21 environments during the cropping seasons of 2009/2010 and 2010/2011. Candidate cultivars with high yield and agronomic stability in the target population of environments (TPE) will then be registered for commercial use. The TPE of the Upland Rice Breeding Program are within the geographical coordinates 1° North to 17°South and 42°West to 70° West and comprise eight states from the Mid-West (Mato Grosso and Goiás), Northeast (Maranhão and Piaúi) and North (Pará, Rondônia, Roraima, and Tocantins). Further details are presented in Supplementary Table 1. Eighteen locations were sampled in the TPE (Figure 1), where trials were laid out in randomized complete blocks with four replications. Experimental plots had four 5 m rows 0.3 m apart from each other, totaling an area of 6 m^2^, where 60 seeds per meter were sown. Seed yield (kg ha^-1^) was measured in the two central rows. Management practices followed the technical recommendations adopted for upland rice in these regions.

#### 2.1.2 Soybean dataset

The soybean dataset is composed of 195 purelines evaluated over three cropping seasons (2019/20, 2020/21, and 2021/22) at 13 locations in the state of Mato Grosso do Sul, in the Central-West region of Brazil (Figure 1). Trials were implemented under rainfed conditions and were conducted by the Mato Grosso do Sul Foundation (Fundação MS) in 49 environments. The experimental design was randomized complete blocks with three replications. The plots consisted of five 12 m long rows, spaced at 0.5 m, with a total area of 30 m^2^. Seed yield (kg ha^-1^) was measured in the three central rows and corrected for 13% moisture. Weed and pest control were carried out following the recommendations for the region.

### 2.2 GIS-FA workflow

Here, we will summarize the procedures for applying the GIS-FA methodology. The method was created to evaluate the overall performance and stability of genotypes in untested environments and to plot the spatial prediction on thematic maps. This enables breeders to define strategies for recommending adaptable cultivars, prospect new target environments that maximize genetic gain through selection, and define breeding zones based on the pattern of environmental features. The procedures to apply the GIS-FA are:

- **Step 1 -Geographic data collection from tested and untested environments**: to implement the GIS-FA method, it is imperative to acquire geographic information. This includes but is not limited to latitude and longitude. For the tested environments, such data can be obtained *in situ* in the experimental area or via Geographic Information System (GIS) tools. For the untested environments, one can sample pixels (coordinates) of the breeding region (or the area under consideration for prediction). These pixels must be representative of the different environmental conditions found in the breeding region. We detail the sampling process adopted in this study in section 2.3.
- **Step 2 -Environmental data collection**: information on the sowing and harvest time for each genotype is needed in this step. The process of envirotyping (data collection and processing) is crucial for understanding the environmental drivers of the GxE interaction and how the environment shapes the development of the plant (Cooper et al, 2014; Xu, 2016; Costa-Neto et al, 2021a). Environmental features can be obtained in the form of *in situ* data, or available in raster format (e.g., historical series for a given geographic point). Other methods of obtaining this data include meteorological stations, National Centers for Environmental Information (NCEI) (Noaa, 2023), Climate Forecast System Reanalysis (CFSR) (Ncei, 2018), European Centre for Medium-Range Weather Forecasts (ECMWF) (Ecmwf, 2023), Global Historical Climatology Network (GHCN) (Ghcnd, 2023), NASA Earth Observing System Data and Information System (EOSDIS) (Eosdis, 2023), worldclim (Fick and Hijmans, 2017), Climatologies at high resolution for the Earth’s land surface areas (CHELSA) (Chelsa, 2023), and Climate Research Unit Time-Series (CRU TS). Soil data can be collected through analyses in the experiment itself or in databases, such as SoilGrids (SoilGrids, 2022). We detail the collection of environment features (EF) in both datasets analyzed in section 2.3. The use of environmental features in statistical-genetic models is based on Shelford’s Law (Shelford, 1911), where the growth of the species is regulated by environmental factors (range of maximum and minimum values). In this case, there is an association between the environmental marker and the evaluated genotype. Environmental features can also be used to characterize tested and untested environments, enabling the determination of how similar the sampled points are to the TPE (see section 2.4 for details).
- **Step 3 -Phenotypic data analysis**: In this step, FA models with different numbers of factors are fitted, and we must choose one of them based on parsimony and/or explanatory ability (process detailed in section 2.5.1). After choosing the model, we use the FA selection tools (Stefanova and Buirchell, 2010; Smith and Cullis, 2018) to build a selection index and select the best-ranking genotypes across environments (details in section 2.5.3).
- **Step 4 -Prediction for the untested environments**: The matrix of rotated loadings of the chosen FA model is used to train a PLS regression model with the gathered environmental features. The goal is to predict the factor loading of untested environments only by providing the model with environmental information about these locations. Once the loadings are predicted, they are used in linear combinations with the experimental genotypes’ factor scores to predict the eBLUPs in untested environments. This process is thoroughly detailed in section 2.6.
- **Step 5 -Map-based recommendation**: The prediction phase provides the performance of each genotype in the new locations that were sampled in the first step. To extrapolate to the whole breeding region, an interpolation process is required (detailed in section 2.4). We proposed three types of thematic maps considering the interpolation: (*i*) adaptation zones, which allows for the identification of adaptation areas for each genotype, i.e. areas where genotypes are expected to have better responses to the local’s environmental effects; (*ii*) which-won-where, used to identify the most promising experimental genotypes in the breeding region; and (*iii*) pair-wise comparisons, which compares the performance of two genotypes (or a genotype and a commercial check) in the untested environments. At *i* and *iii*, one can make a pre-selection of which genotypes to evaluate using the FA selection tools, and perform a detailed study about these selection candidates’ adaptation throughout the breeding region.

### 2.3 Environmental information

We used 32 EF in this study: three geographical coordinates (altitude, latitude, and longitude), 16 related to the weather conditions, and 13 soil traits (Table 1). The weather covariates for each environment were obtained as daily averages for the growing season (i.e., between sowing and harvest dates) and processed using the `R` (version 4.2.3, R Core Team, 2023) package `EnvRtype` (Costa-Neto et al, 2021c), which downloads raw data from NASA database (Sparks, 2018; NasaPower, 2022). Most of the soil covariates for each location (i.e., latitude/longitude combination) were acquired using the `geodata` package (Hijmans et al, 2023), which downloads rasters from the Soil-Grids platform (SoilGrids, 2022). Only the rasters of soil temperature, isothermality, temperature seasonality, and mean diurnal range were manually downloaded from the platform of Lembrechts et al (2022). Soil rasters were downloaded for a 5-15 cm depth interval, have a resolution of 30 arc seconds with each pixel representing an area of approximately 1 km^2^, and were processed with the `raster` package (Hijmans, 2020).

**Table 1.** Summary statistics of the 32 environmental features classified into three groups: geographical, climatic, and soil-related. Climatic covariates were obtained from 2000 to 2021.

| Group | Environmental information | ID | Unit | Rice data |  |  | Soybean data |  |  |
| --- | --- | --- | --- | --- | --- | --- | --- | --- | --- |
|  |  |  |  | Min | Mean | Max | Min | Mean | Max |
| Geographical | Altitude | alt | meters (m) | 70.00 | 410.19 | 1033.00 | 234.00 | 410.35 | 661.00 |
|  | Latitude | lat | graus (°) | -16.85 | -11.68 | 2.90 | -23.10 | -21.66 | -19.38 |
|  | Longitude | lon | graus (°) | -60.75 | -52.05 | -42.65 | -56.55 | -54.65 | -52.72 |
| Climatic | All sky insolation incidents on a horizontal surface | sw | MJ/m<sup>2</sup>/day | 16.43 | 18.90 | 20.83 | 21.38 | 22.65 | 24.69 |
|  | Clear sky insolation incident on a horizontal surface | lw | MJ/m<sup>2</sup>/day | 380.72 | 405.20 | 424.89 | 401.17 | 408.72 | 417.74 |
|  | Total precipitation | prec | mm/day | 4.58 | 7.23 | 9.77 | 1.87 | 3.65 | 6.32 |
|  | Relative humidity | rh | % | 79.15 | 85.47 | 91.92 | 57.30 | 66.27 | 77.09 |
|  | Slope of saturation vapour pressure curve | spv | kPa°C | 0.16 | 0.19 | 0.22 | 0.19 | 0.22 | 0.24 |
|  | Potential evapotranspiration | etp | mm.day | 7.53 | 8.64 | 9.36 | 9.88 | 10.51 | 11.47 |
|  | Deficit by precipitation | pept | mm.day | -4.49 | -1.40 | 2.21 | -9.60 | -6.86 | -4.01 |
|  | Vapour pressure deficit | vpd | kPa | 0.36 | 0.60 | 0.96 | 0.94 | 1.70 | 2.22 |
|  | Wind speed at 2 m above ground | ws | m/s | 0.06 | 1.04 | 1.82 | 0.62 | 1.69 | 2.14 |
|  | Dew/Frost Point Temperature | tdew | °C | 18.50 | 22.03 | 24.07 | 17.90 | 19.50 | 21.02 |
|  | Daily temperature range | trange | °C day | 5.65 | 7.22 | 8.87 | 9.36 | 12.16 | 14.00 |
|  | Temperature at 2 m above ground | tmean | °C | 21.99 | 24.84 | 27.40 | 25.06 | 27.48 | 28.96 |
|  | Maximum temperature at 2 m above ground | tmax | °C | 26.69 | 28.69 | 31.71 | 29.94 | 33.91 | 36.08 |
|  | Minimum temperature at 2 m above ground | tmin | °C | 17.82 | 21.47 | 23.58 | 20.53 | 21.75 | 22.80 |
|  | Growing degree days | gdd | °C d<sup>-1</sup> | 14.26 | 17.08 | 19.65 | 17.26 | 19.83 | 21.19 |
|  | Effect of temperature on radiation use efficiency | frue | - | 0.65 | 0.78 | 0.89 | 0.78 | 0.89 | 0.94 |
| Soil | Bulk density of the fine earth fraction | bdod | kg dm<sup>-3</sup> | 1.10 | 1.27 | 1.40 | 1.20 | 1.28 | 1.40 |
|  | Clay (<0.002 mm) in fine earth | clay | % | 18.00 | 28.58 | 42.00 | 16.00 | 36.22 | 52.00 |
|  | Silt (0.002-0.05 mm) in fine earth | silt | % | 11.00 | 19.88 | 32.00 | 10.00 | 17.76 | 23.00 |
|  | Sand (>0.05 mm) in fine earth | sand | % | 39.00 | 51.65 | 69.00 | 25.00 | 45.94 | 66.00 |
|  | Volume fraction of coarse fragments (>2 mm) | cfvo | % | 1.00 | 4.38 | 11.00 | 2.00 | 3.59 | 5.00 |
|  | Nitrogen content | nit | g kg<sup>-1</sup> | 0.80 | 1.48 | 2.40 | 1.30 | 1.69 | 2.10 |
|  | Organic carbon density | ocd | kg m<sup>-3</sup> | 1.90 | 2.46 | 3.20 | 2.00 | 2.45 | 3.00 |
|  | pH (H<sub>2</sub>O) | phh2o | - | 4.30 | 5.27 | 5.80 | 5.20 | 5.39 | 5.80 |
|  | Soil organic carbon in fine earth | soc | g kg<sup>-1</sup> | 8.80 | 18.63 | 35.40 | 14.10 | 17.85 | 24.20 |
|  | Soil temperature | tsoil | °C | 226.17 | 253.46 | 292.83 | 237.17 | 257.94 | 270.00 |
|  | Temperature seasonality | sts | °C | 86.10 | 155.20 | 255.70 | 242.70 | 349.49 | 401.90 |
|  | Isothermality | iso | - | -84.60 | 13.67 | 30.70 | 15.30 | 19.74 | 23.30 |
|  | Mean diurnal range | mdr | - | -2.00 | 1.17 | 2.40 | 1.90 | 2.47 | 3.00 |

In this study, we aimed to perform spatial predictions using environmental information in a three-step procedure as follows: (*i*) define the scope of the prediction area based on the political borders of the Brazilian states where trials were conducted; (*ii*) perform a sampling approach to generate a cloud of geographical points (latitude/-longitude) in which the environmental data was collected. 50 points were randomly sampled from each municipality within states, ensuring an unbiased sampling of possible environmental conditions in the states; and (*iii*) using data from *ii*, perform a spatial interpolation to cover the area of the entire state(s) and compute the spatial predictions. In *ii*, the soil-related environmental features were obtained as previously described for the tested environments, and for the weather environmental features, monthly averages were obtained from 2000 to 2021. Further details are given in the next sections.

### 2.4 Environmental similarity and interpolation grid

The Euclidean distances between the observed and unobserved (i.e., sampled points) environments were used to quantify the environmental similarity using the package `pdist` (Wong, 2022). Let **W** be a *J* × *P* matrix of scaled values of *P* environmental features with the *J* observed environments, and let **Ω** be the matrix with the same information but for the *U* unobserved environments. Then, the Euclidean distance between an observed environment *j* and an unobserved environment *u* (*D*_ju_) is given by the distances between the rows of **W** and **Ω** that correspond to *j* and *u*, respectively:

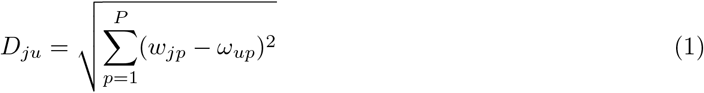

where *w*_jp_ and *ω*_up_ are entries of **W** and **Ω** that contain the value of the *p*^th^ EF for the *j*^th^ tested environment and *u*^th^ untested environment, respectively.

After calculating the distance between all *J* and *U* environments, we expanded these results for all possible environments in the delimited prediction area using the inverse distance weighting (IDW) interpolation method. The IDW was performed using the spatstat package (Baddeley et al, 2015). Let *u*^⋆^ represent an untested and unsampled environment (*u*^⋆^ = 1, 2, … , *U* ^⋆^, with *U* ^⋆^ *>> U*). The Euclidean distance between a given *j* and *u*^⋆^ is defined as:

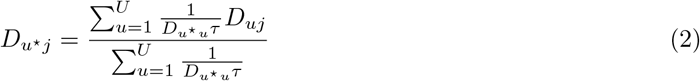

where *D*_*u⋆ u*_ is the Euclidean distance between *u*^⋆^ and *u*, and *τ* is a weight defined by cross-validation (CV). Values ranging from 0.1 to 5.0 by 0.1 were tested in the CV, and the value that provided the lowest mean squared error between the predicted and observed values in the sampled points was adopted.

Once we have performed the interpolation and obtained the Euclidean distances between all tested and untested environments, we considered that the environmental similarity between the *u*^th^ (or *u*^*\**th^) untested environment and the *j*^th^ tested environment is the minimum distance of *u* (or *u*^⋆^) to any *j*:

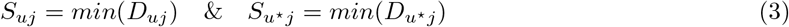

### 2.5 Phenotypic analysis

The phenotypic analyses across environments for both data sets were performed with the following linear mixed model (Henderson, 1949, 1950) using the ASReml-R package (version 4.1.2, Butler, 2021). Variance components were estimated by residual maximum likelihood (Patterson and Thompson, 1971).

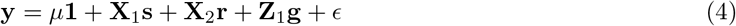

where **y** is the vector of phenotypic records, *µ***1** is the intercept; **s** is the vector of fixed effects of environments with design matrix **X**_1_; **r** is the fixed vector of within-environments block effects with design matrix **X**_2_; **g** is the vector of random genotypic effects nested within environments with incidence matrix **Z**_1_; and *ϵ* is the residual term. The distributional assumptions of **g**, and for *ϵ* are detailed below.

Using the available information about the coordinates (row and column) of each plot in the soybean dataset, we implemented a strategy to control the spatial trends in a single-step, following the approach proposed by Gogel et al (2018). In summary, we conducted model testing in each environment, considering spatial analysis. These adjustments included incorporating autoregressive processes in the error term, as well as incorporating linear and non-linear effects as fixed or random terms, as previously demonstrated by Gilmour et al (1997). We identified the best-fitting model for each specific environment. Once we determined the optimal model for each environment, we incorporated the factors from these models into Equation 4. Each added factor followed a block diagonal covariance structure, with non-nil effects only for environments where these factors were presented in the best within-environment model. Detailed information about this procedure is in Supplementary Table 2. For spatial-adjusted trials, the residual effects are distributed as 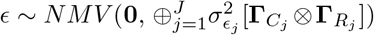, where 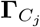 and 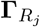 are *C*_j_ × *C*_j_ and *R*_j_ × *R*_j_ autocorrelation matrices, respectively, with *C*_j_ and *R*_j_ being the number of columns and rows in the *j*^th^ trial, respectively. These matrices have 1 on the diagonal and the autocorrelation coefficients that quantify the spatial trends (on column or row directions) on the off-diagonal. For the environments where no spatial adjustment was necessary, 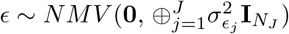, where 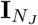 is an identity matrix of order *N*_j_, which correspond to the number of phenotypic records per environment. For the rice dataset, since we did not have access to spatial information, 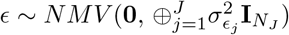.

Genotypic effects were modeled using the factor-analytic covariance structure (Piepho, 1997; Smith et al, 2001):

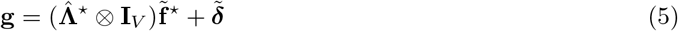

where 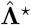 is the *J* × *K* matrix of *K* rotated loadings of the *J* environments 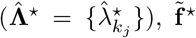 is a vector of *K* rotated scores for the *V* genotypes 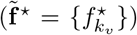, and 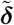 is the vector of the *V J* lack of fit effects 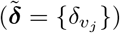. The rotation followed the process described by Smith and Cullis (2018). Under these conditions, 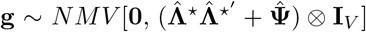, where 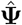 is a *J* × *J* diagonal matrix of environment-wise variances that were not captured by any factor 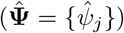, and **I**_V_ is an identity matrix of order *V* .

#### 2.5.1 FA model selection

FA models with different numbers of factors were fitted (FA1 to FA4) and compared in regard to their parsimony and explanatory ability. The parsimony was assessed through the Akaike Information Criterion (AIC, Akaike, 1974), and the explanatory ability was quantified by the Average Semivariances Ratio (ASR, Piepho, 2019; Chaves et al, 2023a). By taking the ratio between the average semivariance of 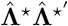 and the average semivariance of 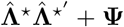, it is possible to investigate the amount of the total covariance that is being captured by the factors of the FA model. The ASR is given as follows:

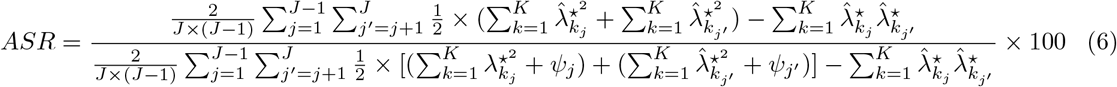

We first defined an ad-hoc threshold of 75% for the explanatory ability. Then, we selected the model with the lowest AIC amongst the ones that surpassed the referred threshold. As complementary information, we also estimated the proportion of genetic variance explained by each factor in each environment (*v*_k_*j*, Smith et al, 2015):

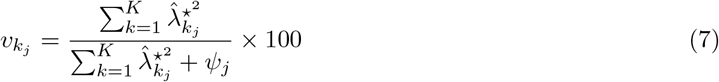

From the best-fit model, we estimated some useful parameters for investigating the experimental precision, such as the environment-wise generalized heritabilities (Cullis et al, 2006) and coefficients of experimental variation (CV), given by the following equations, respectively:

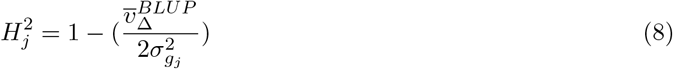

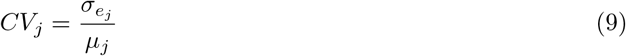

where 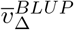 is the average pairwise prediction error variance, 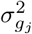 is the genotypic variance for the *j*^th^ environment, taken from the diagonal elements of 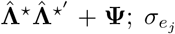 is the estimated residual standard deviation for the *j*^th^ environment, and *µ*_j_ is the mean of the trait for the *j*^th^ environment.

#### 2.5.2 Genotype-by-environment interaction investigation tools

We investigated the GEI dynamics in the datasets through the pairwise genetic correlations between environments, and the partition of the GEI variance into crossover and non-crossover patterns. The pairwise genetic correlation between environments (*ρ*_jj_*′*) is given as follows (Cullis et al, 2010):

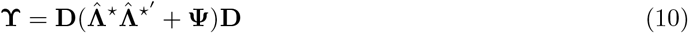

where **ϒ** is a *J* × *J* matrix of genetic correlations, and **D** is a diagonal matrix whose elements are the inverse of the square roots of the diagonal values of 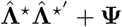.

The decomposition of the GEI variance was performed using the following equation, adapted from Cooper and Delacy (1994):

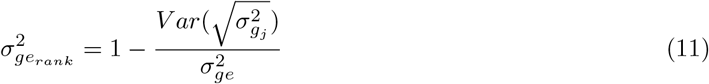

where 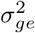 is the variance due to the GEI, obtained by fitting a compound symmetry model. This model has the same structure as Eq. 4, but the variance-covariance matrix of genetic effects has the form 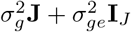, where **J** is an *J* × *J* matrix of ones.

#### 2.5.3 Selection tools

Using the best-fit FA model, we estimated metrics to assess the genotypes’ performance and stability. The performance was measured by the Overall Performance metric (*OP*_v_), obtained as follows (Stefanova and Buirchell, 2010; Smith and Cullis, 2018):

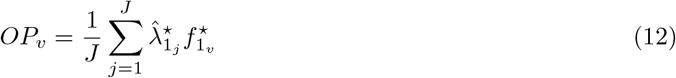

Note that only the first factor is used to compute the *OP*_v_. This factor captures the most significant portion of the total variance. Thus, it provides a generalized measure of the genetic main effects (Supplementary Figure 2 Stefanova and Buirchell, 2010). According to empirical observations of (Smith and Cullis, 2018), this is valid when most of the loadings in the first factor are positive, meaning that there is no (or negligible) crossover GEI in the first factor. Using this principle, the other factors are used to represent stability. Considering that the genetic effect of a given genotype *v* at the *j*^th^ environment, disregarding the lack of fit effect, is 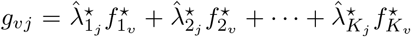, which is equivalent to 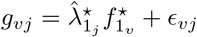, the stability of *v* is given by:

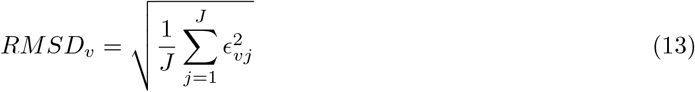

in which *RMSD*_v_ is the root-mean-square deviation of *v*, representing the distance between the point and the slope in a latent regression given by 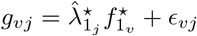 (Smith and Cullis, 2018).

A desirable genotype *i* has high *OP*_i_ and low *RMSD*_i_. Following these principles, we applied a selection index (*SI*_v_) with these metrics (Chaves et al, 2023b; Cowling et al, 2023), given as follows:

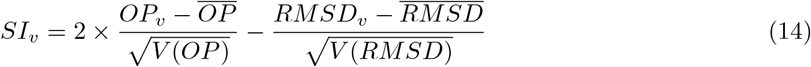

In addition to the selection index, the reliability of the *v*^th^ genotype (Mrode, 2014) was calculated as follows:

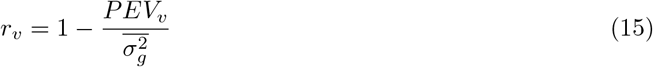

where *PEV*_v_ represents the prediction error variance of the *v*^th^ genotype, and 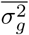 is the average genotypic variance across environments. The reliability metric associated with the selection index is useful to increase the accuracy of selection given both data sets are unbalanced. We adopted a selection intensity of 15% for both datasets.

### 2.6 Spatial predictions in the breeding zone

In this study, GIS tools were used to: (1) collect georeferenced data from the evaluated trials, (2) build environmental markers, and (3) perform spatial predictions for a larger area. Here we used PLS regression (Wold, 1966; Aastveit and Martens, 1986) to perform the predictions. This method is useful when the number of predictors is much larger than the number of observations, and when these predictors are correlated. When PLS is used for predicting genotypic performances in untested environments, the response variable is the genotypic performance in the testing set. In this situation, the response variable is a *J* × 1 vector (**y**) of phenotypic records, if a genotype-wise PLS model is fitted; or a *J* × *V* matrix (**Y**) when a multivariate PLS model is fitted considering all genotypes at once (Monteverde et al, 2019; Costa-Neto et al, 2022). We refer to the multivariate model as GIS-GGE.

We modified GIS-GGE by using the rotated loadings of the tested environments 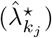 as response variables, instead of the within-environment phenotypic records of the genotypes. We obtained these loadings from the FA model previously chosen (Section 2.5.1). With the predicted loadings and the previously estimated scores for each genotype from the FA model, we can predict the empirical BLUPs of the genotypes in untested environments. The PLS regression model trained with the rotated loadings and environmental features of the tested environments was:

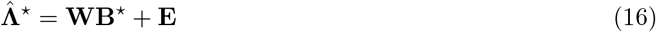

where **B**^⋆^ is a *P* × *K* vector of coefficients, **E** is a *J* × *K* matrix of lack of fit effects, and 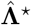 and **W** were previously described (Sections 2.5 and 2.4, respectively). We obtained **B**^⋆^ using a kernel PLS algorithm (Lindgren et al, 1993; Dayal and MacGregor, 1997) implemented in the pls package (Liland et al, 2022). This algorithm is detailed in the Appendix A.

After training the model, we substituted **W** by **Ω** to predict the *K* loadings of the *U* untested environments:

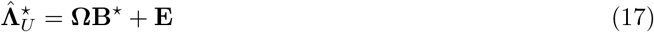

recall from Section 2.3 that **Ω** was built using historical weather data from 2000 to 2021, as well as soil environmental features. Once we predicted the loadings of untested environments, we used them in linear combinations with the previously predicted scores of each genotype (see Section 2.5) to estimate their eBLUPs within untested environments:

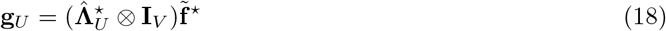

A cross-validation (CV) process is required to obtain **B**^⋆^. We employed a leave-one-out scheme, where data from a single environment was removed (testing set) and predicted using the information provided by the remaining environments (training set). The predicted eBLUPs were then correlated with the actual eBLUPs and eBLUEs of each environment to obtain the predictive ability of the PLS regression model. The model with the number of components with the highest predictive ability was chosen. We leveraged the same CV scheme to compare the predictive ability of GIS-FA and GIS-GGE. In this study, the PLS regression of GIS-GGE was trained with the within-environments empirical eBLUPs of each genotype as response variables.

#### 2.6.1 Thematic maps

Thematic maps combine cartographic principles and GIS tools to represent and analyze spatial geographic phenomena. The incorporation of spatial interpolation methods enables the estimation of values in untested locations, resulting in a seamless representation of the phenomenon (Dabrowski et al, 2021). This facilitates the identification of patterns and trends, providing the decision-making across various fields of study(Costa-Neto et al, 2020).

Recall that **Ω** has *U* rows, and the predictions must be extrapolated to all *U* ^⋆^ untested environments of the targeted area. For this purpose, we used an interpolation process similar to what was described in the Section 2.4. Once the spatial prediction was interpolated in the whole breeding region, we built thematic maps to aid in the visualization and interpretation of results. We built maps with three themes:

- Adaptation zones: These maps depict the expected spatial prediction of each selection candidate across the breeding zone. The adaptation of a genotype to an environment is assessed by the expected response of this genotype when planted in this environment. Thus, in this context, “adaptation” is used as a synonym of specific performance. For better visualization, we divided the predicted eBLUPs into eight categories (from expected yield lower than 2500 kg ha^-1^, to expected yield higher than 4000 kg ha^-1^), and each category was assigned a color.
- Pairwise comparisons: These maps allow for a direct comparison of the expected response of different genotypes in specific environments. Two distinct colors -one for each candidate -were used to indicate which selection candidate was superior in each location on the map. This visual representation helps to quickly identify which selection candidate outperforms the other in each pixel, facilitating the interpretation of competitive advantages among genotypes in specific enviroments.
- Which-won-where: The genotype that achieved the best performance in each location on the map is highlighted. This map provides a clear depiction of the winning genotype for each specific location, enabling a comprehensive understanding of the distribution of top-performing genotypes across the breeding zone.

These maps, just like all the other plots, were built using `ggplot2` package (Wickham, 2016), with add-ins of the packages `ggspatial` (Dunnington, 2023) and `sf` (Pebesma and Bivand, 2023). The shapefiles we used are freely available at the Brazilian Institute of Geography and Statistics (IBGE in the Portuguese acronym) website (https://www.ibge.gov.br/geociencias/organizacao-do-territorio/malhas-territoriais/15774-malhas.html), or can be downloaded using the `geodata` package.

## 3 Results

### 3.1 Experimental accuracy

In the rice dataset, *CV*_j_ ranged from 0.11 (E20) to 0.34 (E13), and 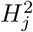 ranged from 0.31 (E08) to 0.78 (E18) (Figure 2a). In the soybean dataset, *CV*_j_ ranged from 0.04 (E31) to 0.17 (E42), and 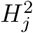 ranges from 0.31 (E18) to 0.77 (E31)(Figure 2b). Spatial trends were modeled in 37 out of 49 soybean trials (Supplementary Table 2).

**Fig. 2.**
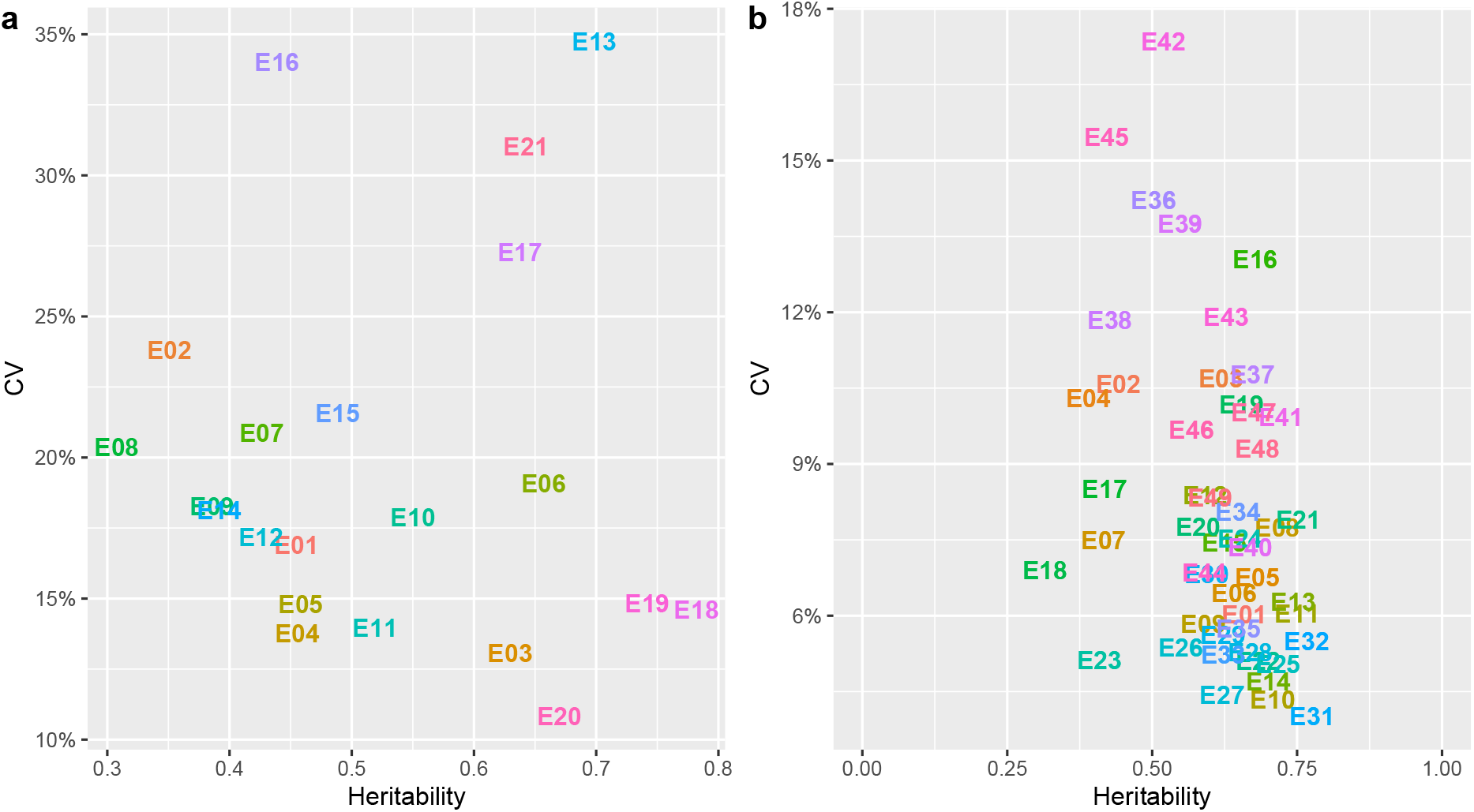
Scatter plot representing the experimental coefficient of variation (CV, on a decimal scale) in the *y* -axis and the generalized heritability in the *x* -axis for grain yield (kg ha^*−*1^) of rice (a) and seed yield (kg ha^−1^) of soybean (b) trials.

### 3.2 Genotype recommendation for tested environments

The FA model with four factors (FA4) met our criteria for both datasets, i.e., it had the lowest AIC among the tested models and more than 75% of explanatory ability (Table 2). This model captured most of the within-environment variance in both datasets (Supplementary Figure 3).

**Table 2.**
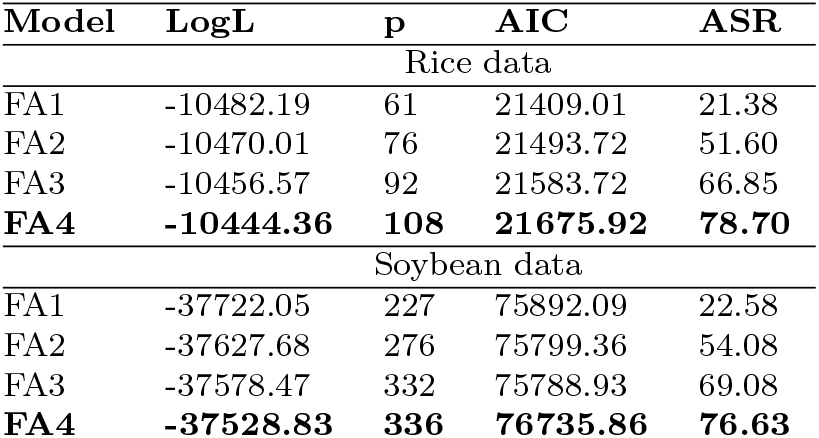
Fitted factor-analytic mixed models for each dataset (rice and soybean) and their respective logarithm of the likelihood function (LogL), number of parameters (p), Akaike information criterion (AIC) and average semivariances ratio (ASR). The selected models are in bold.

| Model | LogL | p | AIC | ASR |
| --- | --- | --- | --- | --- |
| Rice data |  |  |  |  |
| FA1 | -10482.19 | 61 | 21409.01 | 21.38 |
| FA2 | -10470.01 | 76 | 21493.72 | 51.60 |
| FA3 | -10456.57 | 92 | 21583.72 | 66.85 |
| **FA4** | **-10444.36** | **108** | **21675.92** | **78.70** |
| Soybean data |  |  |  |  |
| FA1 | -37722.05 | 227 | 75892.09 | 22.58 |
| FA2 | -37627.68 | 276 | 75799.36 | 54.08 |
| FA3 | -37578.47 | 332 | 75788.93 | 69.08 |
| **FA4** | **-37528.83** | **336** | **76735.86** | **76.63** |

The genotypic correlations ranged from -0.0031 (E07 *vs* E19) to 0.8936 (E13 *vs* E19) for the rice dataset (Figure 3a), and -0.0010 (E07 *vs* E41) to 0.9753 (E031 *vs* E32) for the soybean dataset (Figure 3b). In the rice dataset, environments E17 and E18 exhibited the most contrasting patterns compared to the other environments. Their correlations with the remaining environments were predominantly negative or close to zero. Similarly, in the soybean dataset, negative or negligible correlations were observed for contrasts involving environments E18, E33, E34, E43, E46, and E47. These findings indicate substantial differences between these specific environments and the rest of the dataset. The wide range of correlation magnitudes is reflected by the percentage of crossover GEI in the datasets: 76% and 81% of the total GEI were due to crossover interactions in the rice and soybean datasets, respectively.

**Fig. 3.**
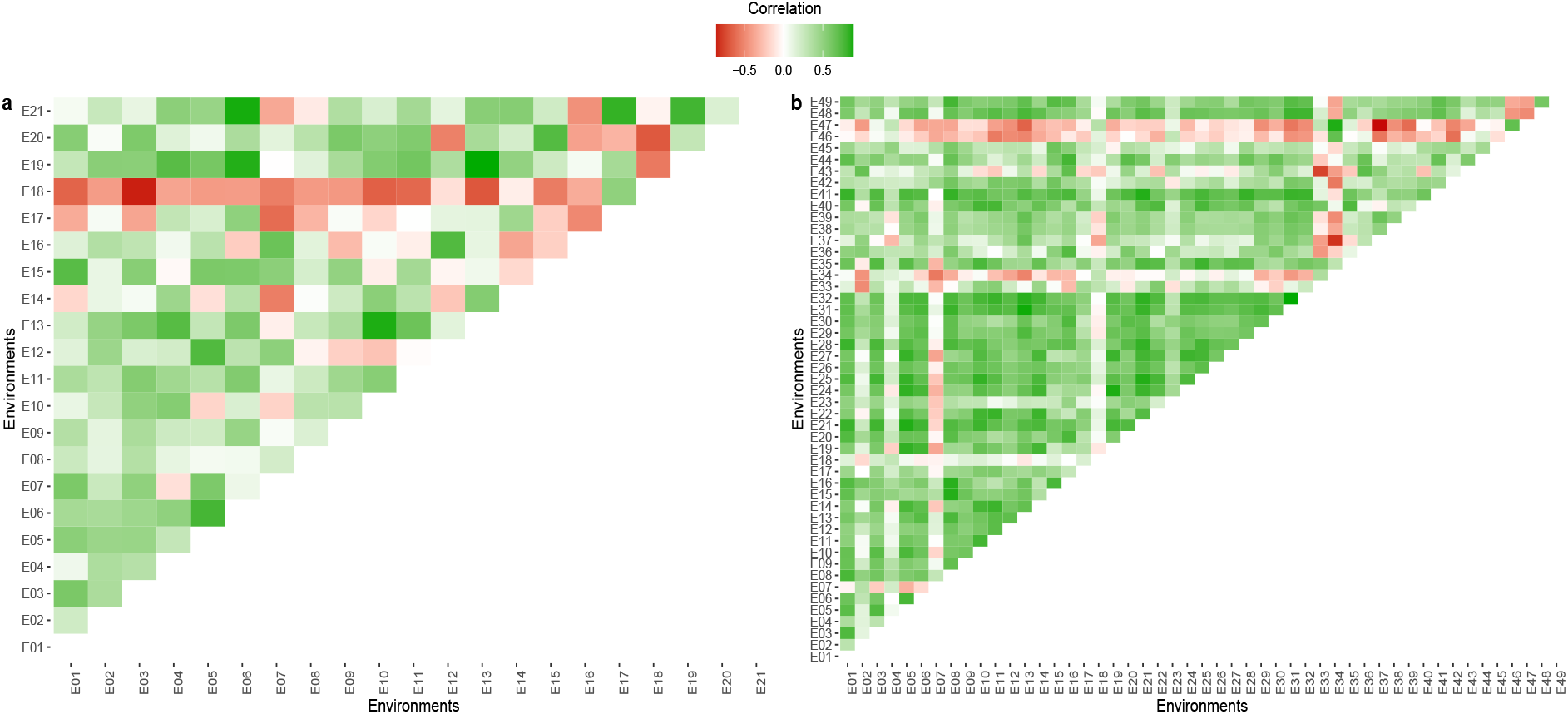
Heatmap representing the genetic correlation between pairs of environments in the rice (a) and soybean (b) datasets. The colour gradient depicts the direction of the correlation: red designates a negative correlation, whereas green represents a positive correlation.

The selected candidates based on the selection index are highlighted in Figure 4. For the rice dataset, in spite of the low reliability, genotypes G23, G18, G29, G31, and G26 stand out due to their high stability. Genotypes G10, G09, G03, and G01 presented high *OP* and reliability. The check treatment (C83) had the highest *OP*, but low stability and reliability if compared to the other selected genotypes (Figure 4a). For soybean, G178, G031, G101, G052, and G035 were the most stable genotypes, G177, G100, G144, G088, and G016 stood out for their high OP. Genotype G16 showed high OP, stability, and reliability (Figure 4b). The reliability of the selected candidates was higher in the soybean dataset, likely due to the higher correlation between environments in the soybean dataset compared to the rice dataset.

**Fig. 4.**
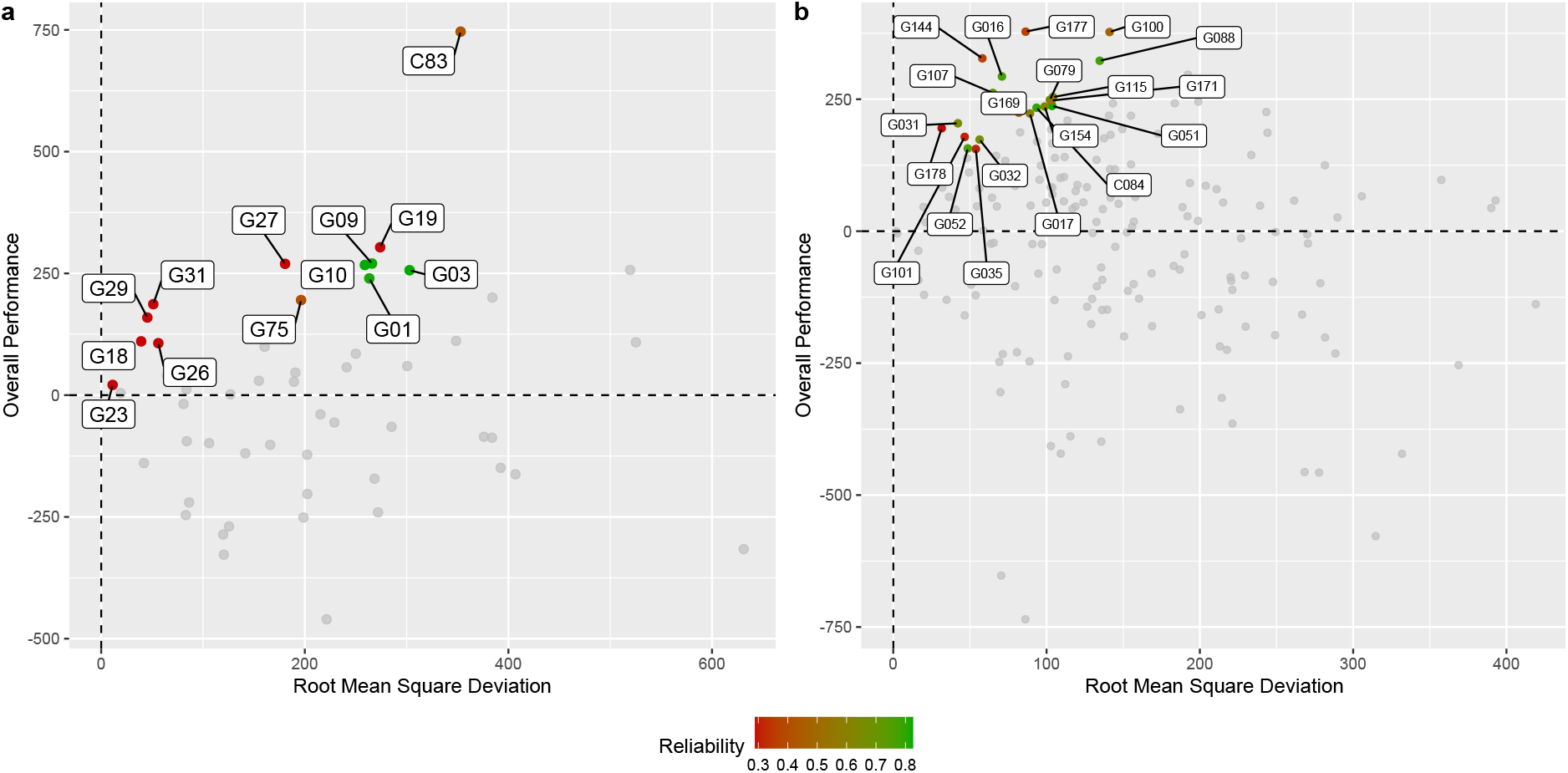
Overall performance (*y* -axis) and root-mean-square deviation (*x* -axis) of the experimental genotypes in the rice (**a**) and soybean (**b**) datasets. The most productive genotypes are oriented towards the upper part in the *y* -axis, and the most stable ones are towards the left in the *x* -axis.

### 3.3 Predictions using environmental markers in untested environments

#### 3.3.1 Environmental similarity

The rice trials are spread all over the breeding region and effectively sample the environmental conditions of the referred area (Figure 5a). On the other hand, the trials in the soybean dataset are concentrated in the central part of the State, and there is an area to the west that has low similarity. This area corresponds exactly to the Pantanal biome, a protected area with legal restrictions to soybean planting (Figure 5b). This is probably the reason why there is no trial in this region.

**Fig. 5.**
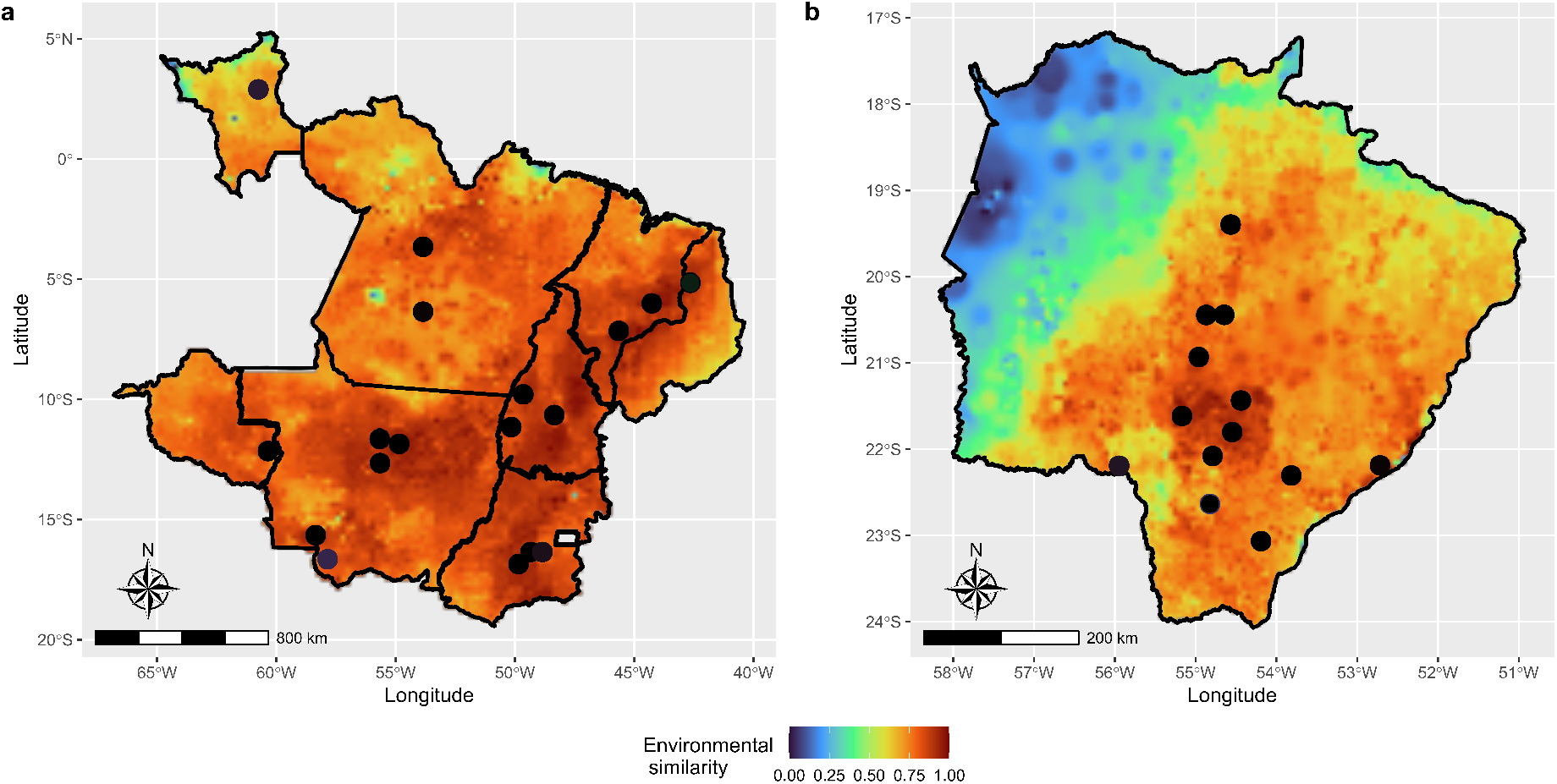
Environmental similarity between tested and untested environments in the target population of environments in the rice (**a**) dataset and in the soybean (**b**) dataset. The warmer the colour, the higher the similarity, and consequently, the higher the prediction reliability. Coloured circles represent the trials’ locations.

#### 3.3.2 GIS-FA validation

In comparison to GIS-GGE, our proposal yields a higher prediction accuracy (simple correlation between predicted and observed values) for both datasets. For predicting eBLUEs, GIS-FA is 10% and 1% better than GIS-GGE, in the rice and soybean datasets, respectively. For predicting eBLUPs, GIS-FA is 9% and 5% more effective than GIS-GGE. A second way to assess the predictive ability of the methods is to check the coincidence between the top 10% observed and predicted values (Figure 6). GIS-FA provides more assertive results (Figures 6a and 6b) than GIS-GGE (Figures 6c and 6d). In other words, upon recommending elite candidates based on predicted values, it is more probable that the true top performers are recommended using GIS-FA, than using GIS-GGE. In the rice dataset (Figures 6a and 6c), GIS-FA has an accuracy 13.15 pp higher than GIS-GGE, and in the soybean dataset (Figures 6b and 6d), GIS-FA is 21.19 pp more advantageous than GIS-GGE.

**Fig. 6.**
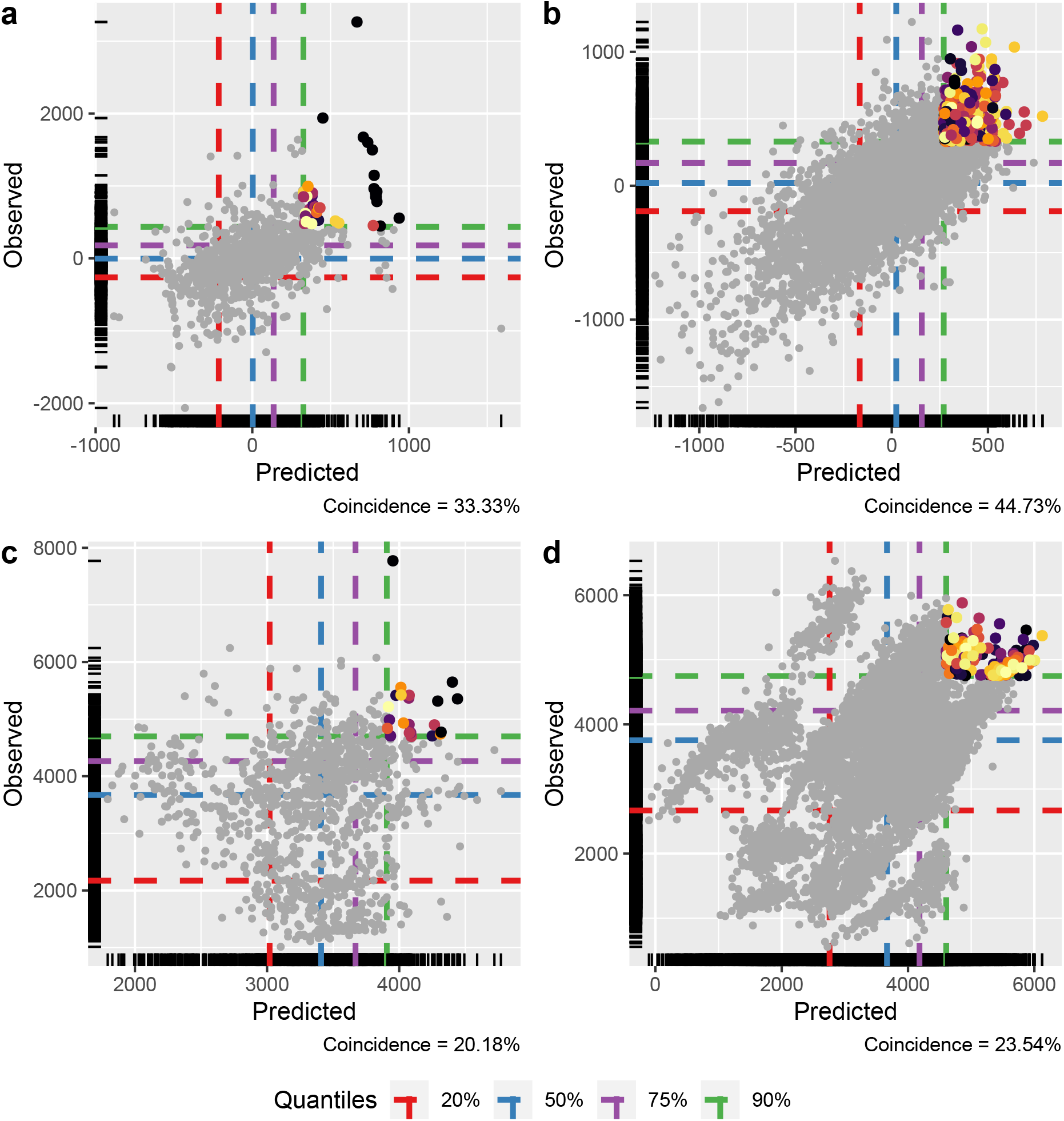
Scatter plot of all predicted values (*x* -axis) in the leave-one-out cross-validation scheme against observed values (*y*-axis). The dashed lines represent the empirical percentiles (20, 50, 75, and 90%) associated with the trait value. The coloured dots represent the coincident selection candidates when selecting the top 10% performers using observed and predicted values. “Coincidence” in the lower left corner of each plot depicts the accuracy of selecting the top 10% using the predicted values. Figures **a** and **b** illustrate the results for the GIS-FA method in the rice and soybean datasets, respectively. Figures **c** and **d** represent the results for the GIS-GGE method in the rice and soybean datasets, respectively.

#### 3.3.3 Thematic maps of adaptation zones

The spatial prediction done by GIS-FA was useful to assess the expected performance of the experimental genotypes within untested environments. This aids in the definition of adaptation zones for each genotype, which are the theme of the maps in Figure 7. For example, G16 of the rice dataset, at Figure 7a, seems to be well adapted only in a small portion of Goiás State (green region), and it responds poorly to the environmental effects of other locations in the breeding region. Conversely, G27 of the rice dataset, at Figure 7b, has a broader spectrum in terms of adaptation in the breeding region. The same interpretation goes for the genotypes of the soybean dataset: G064 (Figure 7c) is an unstable candidate, with a very restricted area where it is better adapted (at the Northern part of the breeding region); and G088 (Figure 7d) is a stable genotype, i.e., it holds alleles that respond favorably to the environmental effects of different location across the State. In each map, we provide *OP* and *RMSD* of the corresponding genotype. We have deliberately chosen two promising candidates (Rice’s G27 and soybean’s G088, which are among the selected in Figure 4), and two low-yielding genotypes (Rice’s G16 and soybean’s G064) to compose Figure 7. Nevertheless, we recommend using *OP* and *RMSD* as criteria to choose the genotype whose adaptation map should be made.

**Fig. 7.**
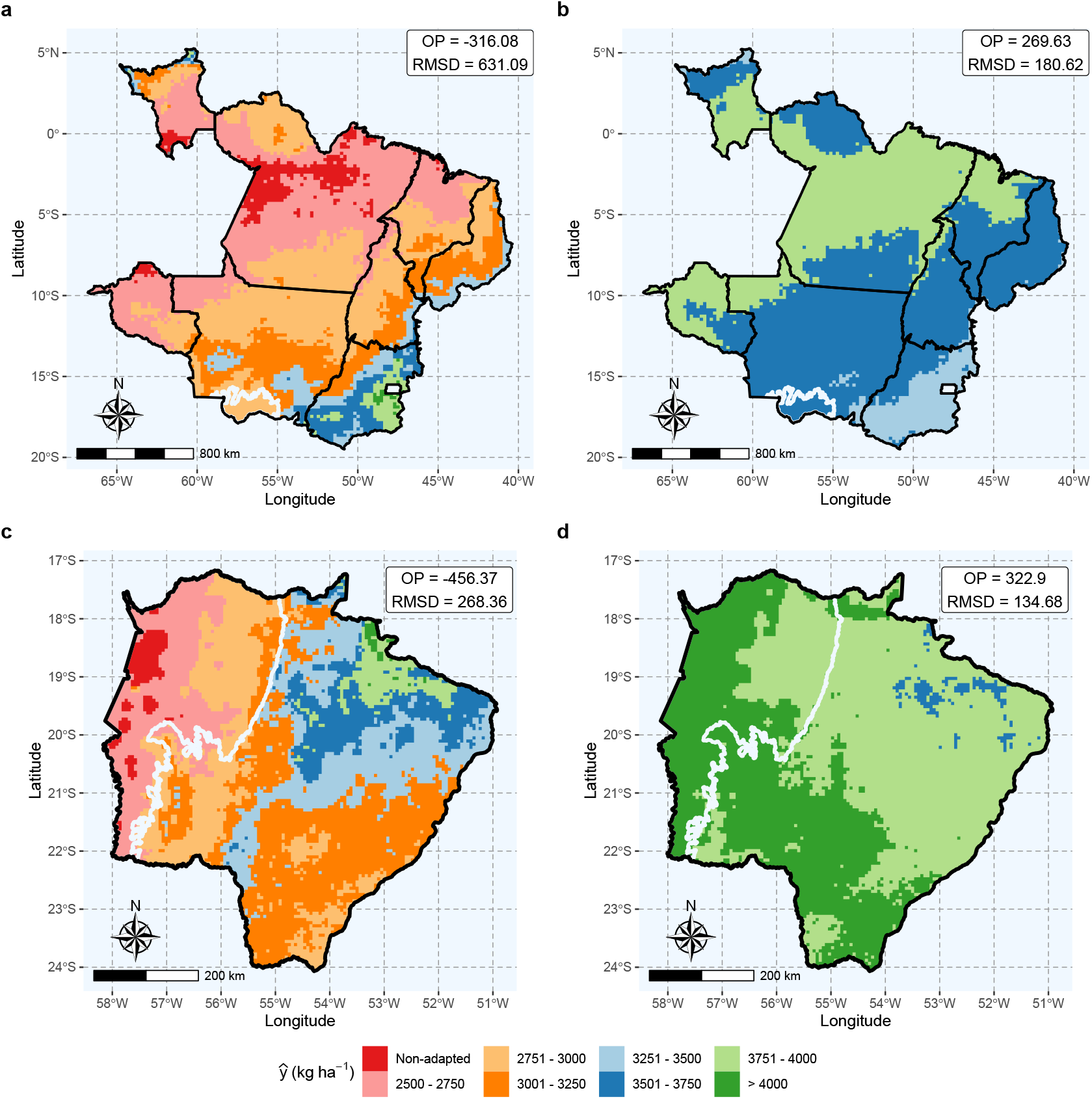
Genotype-wise adaptation map showing the adaptation zones of the genotypes G16 (rice dataset, **a**), G27 (rice dataset, **b**), G064 (soybean dataset, **c**) and G088 (soybean dataset, **d**). The color scale represents the expected yield classes, from non-adapted (intense red) to more than 4000 *kg ha*^−1^ (intense green). The white contour delimits the Pantanal biome. On the upper right of each map, we provide the overall performance (OP) and root-mean-square deviation (RMSD) of each genotype.

#### 3.3.4 Thematic maps of pairwise comparison

To support the decision-making process, we develop a second thematic map: the pairwise comparison maps (Figure 8), which facilitates the comparison of two candidates. Take, for example, G10 and G19 at Figure 4a, and G100 and G177 at Figure 4b. These candidates have a somewhat similar performance according to their *OP* and *RMSD*. However, they are clearly adapted to different zones in the breeding region: G10 have better responses at lower latitudes, and G19 is more suitable to be planted in higher latitudes (Figure 8b); G100 is more adapted to the central region of the soybean’s breeding region, and G177 have better fitness with the environmental conditions of the breeding region’s horizontal extremes (Figure 8d).

**Fig. 8.**
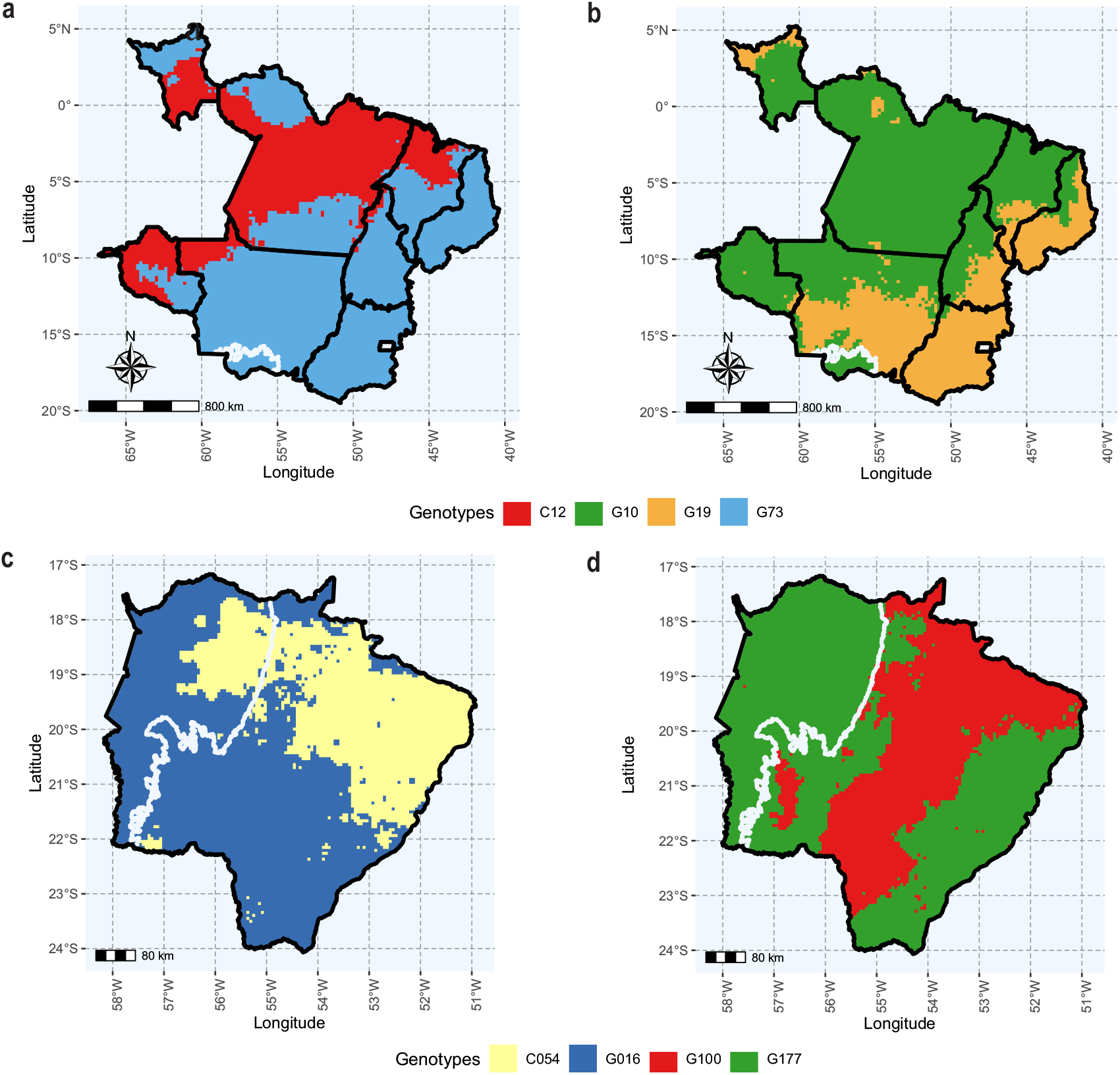
Pairwise comparison map showing the regions within the rice (**a** and **b**) and soybean (**c** and **d**) target population of environments where a selection candidate outperforms a given peer. The colours across the map represent the winning genotype. **a** and **c** are examples of pairwise comparisons between an experimental genotype and a commercial check, whilst **b** and **d** contrast the performance of two promising experimental genotypes along the breeding region. The white contour delimits the Pantanal biome.

#### 3.3.5 Thematic maps of which-won-where

The which-won-where map (Figure 9) shows which experimental genotype is more suitable for a specific environment across the breeding zone. In the rice dataset (Figure 9a), G10 is the most promising experimental genotype in almost all environments at the central and northern portions of the breeding zone, while G19 prevails at the southern and eastern regions. G09, G16, G17, and G20 are the most suitable in specific environments. The breeding region of the soybean dataset is more divided, with G177, G100, G170, and G088 being the most important experimental genotypes, for they are the winners in the widest area. The other selection candidates, including a cultivar check (C054), are the top performers in only a few, restricted environments (Figure 9b).

**Fig. 9.**
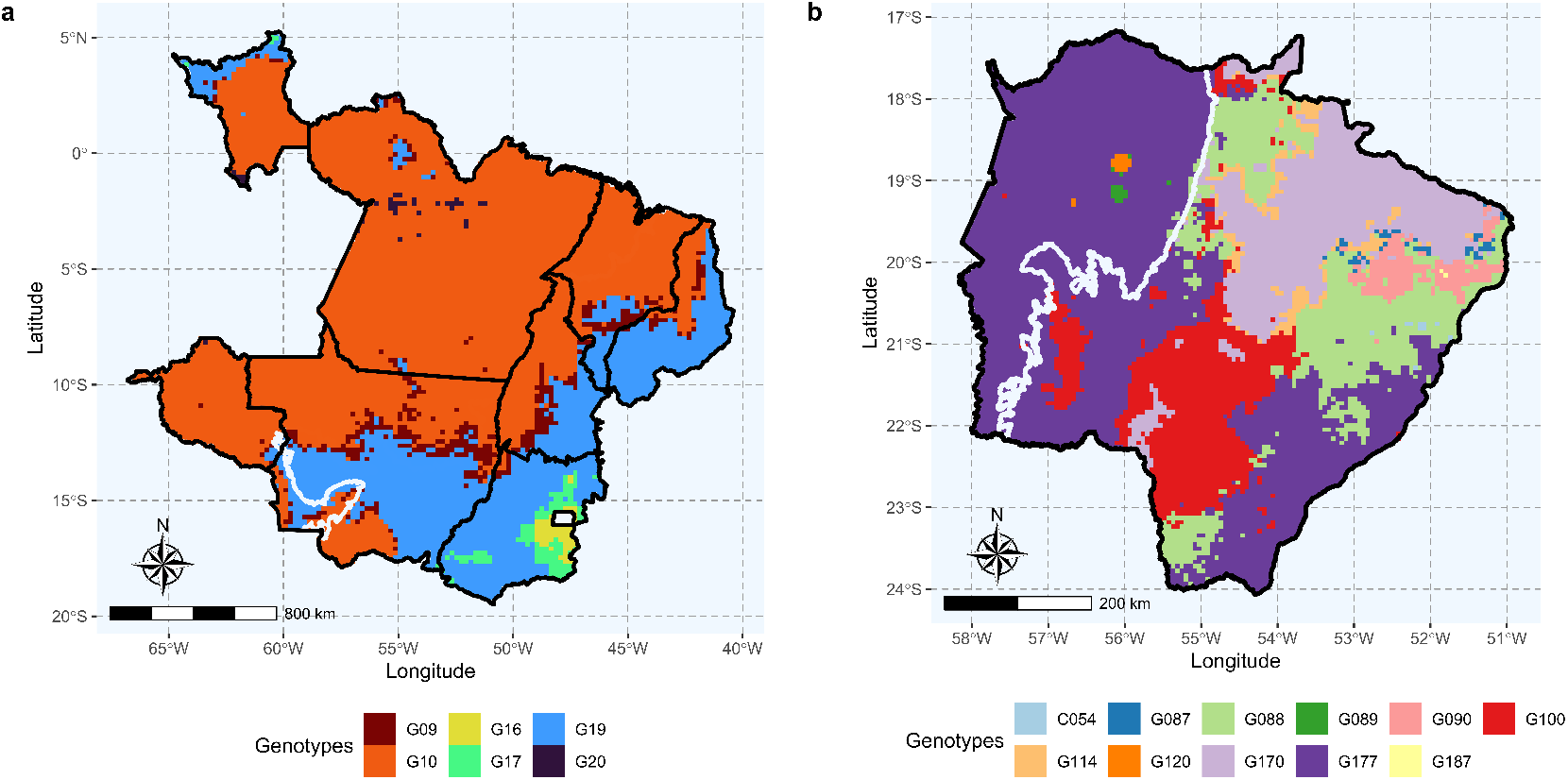
Which-won-where map depicting the most promising genotype at each location across the target population of environments of the rice dataset (**a**) and the soybean dataset (**b**). Each colour represents the experimental genotype that wins in a specific environment within the breeding region. The white contour delimits the Pantanal biome.

## 4 Discussion

The GIS-FA method represents the integration of modern statistical genetics with GIS principles. We showed how GIS-FA can aid plant breeders in decision-making when taking into account the observed performance from observed environments and spatial predictions from untested environments. For observed environments, GIS-FA leverages the resources of FA models to provide useful inferences about the dynamics of the GEI, and to select candidates with high performance and stability using customized selection tools (Stefanova and Buirchell, 2010; Smith and Cullis, 2018). In untested environments, GIS-FA allows for a data-driven recommendation of cultivars, based on spatial predictions according to the soil characteristics, climatic conditions, and parameters from the empirical data (i.e., the factor loadings for genotypes). The GIS-FA method allows for data-driven decision-making with the aid of graphical tools such as the thematic maps: *i*) adaptation zone maps, which depict the expected spatial prediction of each genotype within the entire breeding zone, *ii*) pairwise comparison maps, which are useful to compare the performance of two selection candidates (or a candidate and a commercial check), and the *iii*) which-won-where map, that shows the most promising experimental genotype (the winner) in each location in the breeding zone.

### 4.1 Genotype-by-environment interaction and selection in tested environments

Increasing crop yield and adaptation to different growing conditions are important goals in plant breeding. These traits are the outcomes of a plethora of small quantitative trait loci (QTLs) effects that are highly influenced by the environment (Lynch and Walsh, 1998; Crossa, 2012). In terms of cultivar recommendation in the TPE, the most concerning source of the GEI is the lack of genotypic correlation between environments (Cooper and Delacy, 1994), which was observed in both data sets (Figure 3). As a consequence, it is unlikely the same set of experimental genotypes will have similar performance across uncorrelated environments. In this case, if a global (i.e., across environments) recommendation is needed, metrics such as the selection index that combines performance and stability might be employed. The weight of each metric in the selection index is a breeder’s call (Chaves et al, 2023b). Here, we prioritized performance over stability.

In the GIS-FA method, we leverage the resources of factor-analytic mixed models (Piepho, 1997; Smith et al, 2001) that explore the complexity of the GEI while handling highly unbalanced data sets. Furthermore, FA models allow for a parsimonious estimation of environment-wise genotypic variances and pairwise covariances. These covariances can be used to investigate the dynamics of GEI, as fully described in this study. The efficiency of the GIS-FA method depends on the choice of the number of factors loadings in the FA model, i.e., a poor choice will provide erroneous results. Recall that in GIS-FA, the factor loadings of observed environments are used as the training set, so the loadings of untested environments in the testing set can be predicted. Thus, when selecting the best-fit FA model, selection criteria such as the AIC, and ASR, should always be considered. Naturally, using more factors will provide greater explanatory ability. Nevertheless, it will hinder parsimony and computational efficiency, especially in large data sets.

Assuming that the observed environments faithfully describe the environmental conditions expected all over the breeding zone, the most promising genotypes in the tested environments are probably the best ones in the untested environments. Thereby, the idea is to prioritize selected experimental genotypes when drawing the thematic maps “genotype-wise adaptation” and “pairwise comparisons”.

### 4.2 Spatial interpolations in untested environments

Like molecular markers, EF similarity for either inference or prediction purposes. Inference models aim to determine the effect of each environmental feature on the phenotypic expression and in the GEI effects, which gives a direct analogy of QTL mapping models (Denis, 1988; Van Eeuwijk and Elgersma, 1993; Crossa et al, 1999; Costa-Neto et al, 2021c; Heinemann et al, 2022). In this work, we focused on the environmental-wide predictions, regardless of the particular effect of each EF on the phenotypic expression and GEI. As polygenic models are used to perform whole-genome regressions (Meuwissen et al, 2001), here we assumed that the core of ecophysiological effects captured by the environmental feature could be enough to fit genotype-wise predictions across the spatial grid. The benefits of environmental features in predictive breeding are advantageous in most cases, whether integrated with genomic information or not (de los Campos et al, 2020; Buntaran et al, 2021; Jarquin et al, 2021; Costa-Neto et al, 2022). However, recent work from Crossa et al (2023) demonstrated the simple incorporation of environmental covariates may increase or decrease prediction accuracy depending on the case. Techniques such as feature selection (Crossa et al, 2023) and the exhaustive search (Li et al, 2018) can be considered when selecting environmental features.

#### 4.2.1 Environmental similarity

Environmental similarity maps revealed a need to perform an adequate sampling of the environmental types within a given target breeding region (Figure 5). This entails including samples from various climatic conditions and soil traits that can be encountered in future prediction environments. Essentially, these maps illustrate a metric of reliability of the spatial predictions by benchmarking the similarity among observed environments and the unobserved environments, for they illustrate the environmental similarity between tested and untested environments. In other words, the more similar an untested environment is to a tested environment, the higher the chances of an assertive prediction. The results depicted in the maps of Figure 5 can be attributed to the geographical distribution of trials in relation to the Brazilian biomes [refer to Figure 1 of Chaves et al (2023b) for a map with the Brazilian biomes]. The soybean breeding region comprises two biomes, namely Pantanal (wet low-lands) and Cerrado (highlands savanna conditions). All trials were conducted in Cerrado, which explains the lack of similarity between the TPE and the environments at the Pantanal biome. Consequently, the prediction for this particular region is likely to be compromised. The rice breeding region also includes two biomes, Amazonia (wet tropical rainforest) and Cerrado. Unlike the soybean dataset, there are representative trials from both biomes, providing comprehensive coverage of the relevant environmental conditions.

#### 4.2.2 Predicting using partial least squares regression

The association between partial least square regression, GEI, and environmental features was introduced by Aastveit and Martens (1986) for inference purposes. Their aim was to address challenges related to the curse of dimensionality and multicollinearity in explaining the dynamics of GEI using two datasets. Their model was later expanded to consider the information on molecular markers to investigate QTL-by-environment interaction (Crossa et al, 1999; Vargas et al, 2006). Nevertheless, employing environmental features in statistical models to explain and predict GEI did not reach significant popularity among plant breeders (Vargas et al, 2001; Ortiz et al, 2007; Ramburan et al, 2012; Porker et al, 2020). With the advancement of computational technology and the democratization of “enviromics” resources, PLS became a suitable method for exploring big data to perform spatial predictions of experimental genotypes in new environments (Monteverde et al, 2019; Rincent et al, 2019; Guo et al, 2021; Costa-Neto et al, 2022). In fact, PLS has emerged as a relevant alternative for prediction purposes, even when breeders do not specifically incorporate environmental data into the model (Montesinos-López et al, 2022b,a; Ortiz et al, 2023).

In most studies that employed PLS regression for prediction purposes, the training set typically consisted of the performance *per se* of genotypes and environmental features from the tested environments (Monteverde et al, 2019; Costa-Neto et al, 2022). Our study demonstrated that associating environmental features with the rotated factor loadings of the tested environment yields superior results. Through GIS-FA, we achieved higher prediction accuracy (Table 3) and enhanced the ability to distinguish high-performance experimental genotypes when relying solely on predicted values (Figure 6). By predicting the factor loadings for untested environments, we establish a connection between the observed environmental feature values and the underlying causes of GEI, as well as the genetic covariance that exists between environments. A prior study by Rincent et al (2019) also utilized PLS models to predict latent factors of the AMMI components for untested environments, which enabled them to construct an appropriate covariance structure that improved predictions. The findings of Rincent et al (2019) and the results in this work provide evidence of the potential of using PLS models to indirectly perform spatial predictions by initially predicting the latent elements that contribute to that particular performance. A similar strategy was proposed in a single-step model by Tolhurst et al (2022), who demonstrated the efficiency of combining known and latent environmental features to predict tested and untested environments.

**Table 3.** Prediction accuracy of eBLUEs and eBLUPs using the proposed method GIS-FA, and the conventional method GIS-GGE. For more information about these methods, see the Material and Methods section.

| Model | Prediction | Prediction accuracy |  |
| --- | --- | --- | --- |
|  |  | Rice data | Soybean data |
| GIS-GGE | BLUE | 0.40 | 0.53 |
| GIS-GGE | BLUP | 0.55 | 0.71 |
| GIS-FA | BLUE | 0.44 | 0.55 |
| GIS-FA | BLUP | 0.60 | 0.74 |

#### 4.2.3 Thematic maps

An important feature of GIS-FA is the illustration of the spatial predictions from selection candidates using thematic maps (Figures 7, 8 and 9). Figure 7 offer information on the areas within the breeding zone where the experimental genotypes are expected to thrive. Figure 7 allows the evaluation of the merit of a certain candidate cultivar based on its ability to outperform a commercial cultivar used as a reference or another promising experimental genotype. Figure 9 provides a straightforward solution regarding genotype recommendation across the breeding region, showing which candidate is more suitable for a specific environment across the breeding zone. Thematic maps serve as valuable tools in decision-making, assisting in the allocation of genotypes in the breeding region (Costa-Neto et al, 2020; Bustos-Korts et al, 2022). In addition, the thematic maps offer information on the genotypes’ stability and adaptation from a geographic perspective. Costa-Neto et al (2020) suggested that, in a GIS context, “stability” means lower variability in spatial patterns, while the adaptation regards the expected performance in a specific environment in the breeding region.

One advantage of this approach is the possibility of integrating high-quality satellite images from diverse platforms. Here we used geographic databases freely available on online platforms so that an efficient prediction method without additional cost can be accomplished. Furthermore, implementing partial geographic visualizations can optimize resource allocation when defining the experimental network of trials. The higher resolution of the satellite-based data, it could be to deliver spatial predictions at the farmer’s level. This could benefit the product development and placement stages, by extending this methodology to accommodate satellite-based enviromics also accounting for historical agronomic records.

#### 4.2.4 Future directions

The statistical models of GIS-FA can be improved by integrating molecular information to leverage covariance between relatives and employing more informative EF in the PLS model (Dias et al, 2018; Monteverde et al, 2019; Crossa et al, 2023). The utilization of ecophysiological EF in crop-growth models could enhance our understanding of the link function between the phenotypic expression and environmental factors (Rincent et al, 2019; Costa-Neto et al, 2021a). Other statistical resources - and even artificial intelligence methods -can replace the PLS in the prediction step (Guo et al, 2021; Heinemann et al, 2022). Finally, future research on exploring the potential risks associated with assigning genotypes to specific environments using GIS-FA, possibly through the application of probabilistic methods (Dias et al, 2022), can also be explored.

## Supporting information

GIS-FA Supplementar 1

## Declarations

### Funding

This research was supported by the Minas Gerais State Agency for Research and Development (FAPEMIG), Coordination for the Improvement of Higher Education Personnel (CAPES), and the Brazilian National Council for Scientific and Technological Development (CNPq).

## Conflict of interest

The authors declare no conflict of interest.

## Contributions

M.S.A., S.F.S.C., and K.O.G.D conceived the research. M.S.A. and S.F.S.C. executed the statistical analyses and drafted the initial manuscript. M.D.K. and G.C.N. provided insights into the methodology. L.A.S.D., F.M.F., G.R.P., R.S.A., P.C.S.C., M.D.K., and G.C.N. provided critical revisions of the paper drafts. A.R.G.B. provided knowledge on the structure of the soybean dataset, while A.B.H. and F.B. provided information about the rice dataset. M.S.A., S.F.S.C., and M.D.K. built the tutorial available in the Supplementary Material. All authors approved the final version of the manuscript.

## Data availability

The `R` codes and both datasets used in this study are freely available shortly. The Supplementary Material contains a detailed tutorial with a commented script describing the steps for performing GIS-FA analysis with the soybean dataset.

## Acknowledgments

This work was supported by the Minas Gerais State Agency for Research and Development (FAPEMIG), Brazilian National Council for Scientific and Technological Development (CNPq), Coordination for the Improvement of Higher Education Personnel (CAPES), Mato Grosso do Sul Foundation (Fundação MS), Brazilian Agricultural Research Corporation (Embrapa Rice and Beans), and Federal University of Viçosa (UFV).

## Appendix A Partial least squares regression

Here, we employed the kernel partial least square algorithm (Lindgren et al, 1993; Dayal and MacGregor, 1997) to predict the factor loadings of untested environments. Details about this algorithm are presented below:

Take the following multiple regression as a starting point:

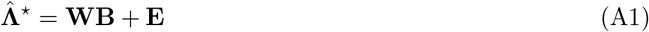

where 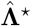 is the *J* × *K* matrix of *K* rotated loadings of the *J* observed environments, **W** is a *J* × *P* matrix of scaled values of *P* environmental features with the *J* observed environments, **B** is a *P* × *K* vector of coefficients, and **E** is a *J* × *K* matrix of lack of fit effects. Note that most of the EF are correlated (Supplementary Figure 4), so **W** has multicollinearity problems, and 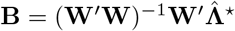 does not yield a proper solution. To overcome this issue, we employed the kernel partial least square regression, which transforms **B** into **B**\*, using the following equation:

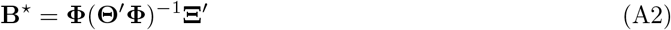

where **Φ** is a *P* × *C* matrix of weights for **W** (**Φ** = {***ϕ***_1_ ***ϕ***_2_ … ***ϕ***C}), with *C* being the number of PLS components; **Θ** is a matrix of loadings for **W** (**Θ** = {***θ***_1_ ***θ***_2_ … ***θ***_C_}) and has the same dimension of **Φ**, and **Ξ** is a *K* × *C* matrix of weights for **Λ** (**Ξ** = {***ξ***_1_ ***ξ***_2_ … ***ξ***C}). We describe the cross-validation procedure that defined the number of components (*c* = 1, 2, … , *C*) in Section 2.6. **Φ, Θ** and **Ξ** were defined using an iterative process that leveraged kernel functions of **W** and **Λ**. First, ***ϕ***c is estimated as the eigenvector that is equivalent to the largest eigenvalue of the kernel 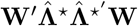. We used this vector to initialize an iterative process, whose number of repetitions is equivalent to *C*. Let **R** = **Φ**(**Θ**^*′*^**Φ**)^−1^, with **R** = {**r**_1_ **r**_2_ … **r**_C_}. In the first iteration, **r**_1_ = ***ϕ***1. Subsequently, 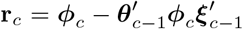. On each iteration, ***θ***_c_ and ***ξ***c are estimated as follows:

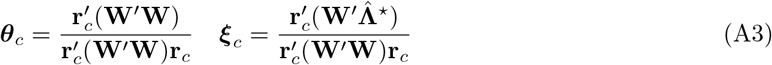

The solutions of these equations are stored in **Θ** and **Ξ**, respectively, and used to update the covariance matrix for the next iteration as follows:

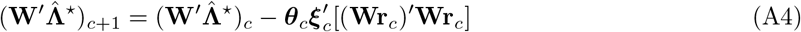

When the iteration process is finished, **B**\* provides a proper solution to Equation A1 and can be used for prediction purposes. We used **B**\* at Equation 16 to train the PLS model, and at Equation 17 to perform the predictions.

## Notes

### Competing Interest Statement

The authors have declared no competing interest.

### Summary of Updates

The new version of the article has inserted citations that complement the ideas of the discussion.

## References

Aastveit AH, Martens H (1986) ANOVA interactions interpreted by partial least squares regression. Biometrics 42(4):829–844. https://doi.org/10.2307/2530697

Akaike H (1974) A new look at the statistical model identification. IEEE Trans Autom Control 19:716–723. https://doi.org/10.1109/TAC.1974.1100705

Alvares CA, Stape JL, Sentelhas PC, et al (2013) Köppen’s climate classification map for brazil. Meteorol Zeitschrift 22:711–728. https://doi.org/10.1127/0941-2948/2013/0507

Annicchiarico P, Bellah F, Chiari T (2006) Repeatable genotype × location interaction and its exploitation by conventional and gis-based cultivar recommendation for durum wheat in algeria. European Journal of Agronomy 24:70–81. https://doi.org/10.1016/j.eja.2005.05.003

Baddeley A, Rubak E, Turner R (2015) Spatial point patterns: methodology and applications with R. Journal of Statistical Software 75:1–6. https://doi.org/10.18637/jss.v075.b02

Bakare MA, Kayondo SI, Aghogho CI, et al (2022) Parsimonious genotype by environment interaction covariance models for cassava Manihot esculenta. Frontiers in Plant Science 13:978248. https://doi.org/10.3389/fpls.2022.978248

Balestre M, Von Pinho RG, Souza JC, et al (2009) Genotypic stability and adaptability in tropical maize based on AMMI and GGE biplot analysis. Genetics and Molecular Research 8(4):1311–1322. https://doi.org/10.4238/vol8-4gmr658

Beebe S, Lynch J, Galwey N, et al (1997) A geographical approach to identify phosphorus-efficient genotypes among landraces and wild ancestors of common bean. Euphytica 95:325–338. https://doi.org/10.1023/A:1003008617829

Buntaran H, Forkman J, Piepho HP (2021) Projecting results of zoned multi-environment trials to new locations using environmental covariates with random coefficient models: accuracy and precision. Theoretical and Applied Genetics 134:1513–1530. https://doi.org/10.1007/s00122-021-03786-2

Bustos-Korts D, Boar MP, Layton J, et al (2022) Identification of environment types and adaptation zones with self-organizing maps: applications to sunflower multi-environment data in europe. Theoretical and Applied Genetics 135:2059–2082. https://doi.org/10.1007/s00122-022-04098-9

Butler D (2021) Asreml: fits the linear mixed model. URL http://www.vsni.co.uk,r package version 4.1.0.160

de los Campos G, Pérez-Rodríguez P, Bogard M, et al (2020) A data-driven simulation platform to predict cultivars’ performances under uncertain weather conditions. Nature Communications 11:4876. https://doi.org/10.1038/s41467-020-18480-y

Chapman S, Barreto H, et al (1996) Using simulation models and spatial databases to improve the efficiency of plant breeding programs. Plant adaptation and crop improvement pp 563–587

Chaves SFS, Alves RS, Dias LAS, et al (2023a) Analysis of repeated measures data through mixed models: An application in Theobroma grandiflorum breeding. Crop Science 63(4):2131–2144. https://doi.org/10.1002/csc2.20995

Chaves SFS, Evangelista JSPC, Trindade RS, et al (2023b) Employing factor analytic tools for selecting high-performance and stable tropical maize hybrids. Crop Science 63(3):1114–1125. https://doi.org/10.1002/csc2.20911

Chelsa (2023) Glimatologies at high resolution for the earth’s land surface areas. URL https://chelsa-climate.org/

Cooper M, Delacy IH (1994) Relationships among analytical methods used to study genotypic variation and genotype-by-environment interaction in plant breeding multi-environment experiments. Theoretical and Applied Genetics 88:561–572. https://doi.org/10.1007/BF01240919

Cooper M, Messina CD (2021) Can we harness “enviromics” to accelerate crop improvement by integrating breeding and agronomy? Front Plant Sci 12:735143. https://doi.org/10.3389/fpls.2021.735143

Cooper M, Messina CD, Podlich D, et al (2014) Predicting the future of plant breeding: complementing empirical evaluation with genetic prediction. Crop and Pasture Science 65:311. https://doi.org/10.1071/CP14007

Cooper M, Messina CD, Tang T, et al (2022) Predicting genotype×environment×management (G×E×M) interactions for the design of crop improvement strategies, pp 467–585. https://doi.org/10.1002/9781119874157.ch8

Coppock JT, Rhind DW (1991) Geographical information systems: principles, techniques, management and applications, Longman Scientific Technical, pp 21–43

Costa-Neto G, Fritsche-Neto R (2021) Enviromics: bridging different sources of data, building one framework. Crop Breeding and Applied Biotechnology 21(spe):e393521S12. https://doi.org/10.1590/1984-70332021v21Sa25

Costa-Neto G, Morais Júnior OP, Heinemann AB, et al (2020) A novel GIS-based tool to reveal spatial trends in reaction norm: upland rice case study. Euphytica 216:37. https://doi.org/10.1007/s10681-020-2573-4

Costa-Neto G, Crossa J, Fritsche-Neto R (2021a) Enviromic assembly increases accuracy and reduces costs of the genomic prediction for yield plasticity in maize. Frontiers in Plant Science 12:717552. https://doi.org/10.3389/fpls.2021.717552

Costa-Neto G, Fritsche-Neto R, Crossa J (2021b) Nonlinear kernels, dominance, and envirotyping data increase the accuracy of genome-based prediction in multi-environment trials. Heredity 126(1):92–106. https://doi.org/10.1038/s41437-020-00353-1

Costa-Neto G, Galli G, Carvalho HF, et al (2021c) EnvRtype: a software to interplay enviromics and quantitative genomics in agriculture. G3 Genes|Genomes|Genetics 11(4):jkab040. https://doi.org/10.1093/g3journal/jkab040

Costa-Neto G, Crespo-Herrera L, Fradgley N, et al (2022) Envirome-wide associations enhance multi-year genome-based prediction of historical wheat breeding data. G3: Genes|Genomes|Genetics 13(2):jkac313. https://doi.org/10.1093/g3journal/jkac313

Cowling WA, Castro-Urrea FA, Stefanova KT, et al (2023) Optimal contribution selection improves the rate of genetic gain in grain yield and yield stability in spring canola in Australia and Canada. Plants 12:383. https://doi.org/10.3390/plants12020383

Crossa J (2012) From genotype × environment interaction to gene × environment interaction. Current Genomics 13(3):225–244. https://doi.org/10.2174/138920212800543066

Crossa J, Vargas M, Van Eeuwijk FA, et al (1999) Interpreting genotype× environment interaction in tropical maize using linked molecular markers and environmental covariables. Theoretical and applied genetics 99:611–625. https://doi.org/10.1007/s001220051276

Crossa J, Yang RC, Cornelius PL (2004) Studying crossover genotype × environment interaction using linear-bilinear models and mixed models. Journal of Agricultural, Biological, and Environmental Statistics 9(3):362–380. https://doi.org/10.1198/108571104x4423

Crossa J, Montesinos-López OA, Crespo Herrera LA, et al (2023) Do feature selection methods for selecting environmental covariables enhance genomic prediction accuracy? Frontiers in Genetics 14:7016. https://doi.org/10.3389/fgene.2023.1209275

Cullis B, Beeck CP, Cowling WA (2010) Analysis of yield and oil from a series of canola breeding trials. part II: exploring vxe using factor analysis. Genome 53:1002–1016. https://doi.org/10.1139/G10-080

Cullis BR, Smith AB, Coombes NE (2006) On the design of early generation variety trials with correlated data. Journal of Agricultural, Biological, and Environmental Statistics 11:381. https://doi.org/10.1198/108571106X154443

Cullis BR, Jefferson P, Thompson R, et al (2014) Factor analytic and reduced animal models for the investigation of additive genotype-by-environment interaction in outcrossing plant species with application to a pinus radiata breed-ing programme. Theoretical and Applied Genetics 127:2193–2210. https://doi.org/10.1007/s00122-014-2373-0

Dabrowski PS, Specht C, Specht M, et al (2021) Three-dimensional thematic map imaging of the yacht port on the example of the polish national sailing Centre Marina in Gdańsk. Applied Sciences 11(15):7016. https://doi.org/10.3390/app11157016

Dayal BS, MacGregor JF (1997) Improved PLS algorithms. Journal of Chemometrics 11(1):73–85. https://doi.org/10.1002/(SICI)1099-128X(199701)11:1⟨73::AID-CEM435⟩3.0.CO;2-#

Denis BJ (1988) Two way analysis using covarites1. Statistics 19(1):123–132. https://doi.org/10.1080/02331888808802080

Dias KOG, Gezan SA, Guimarães CT, et al (2018) Improving accuracies of genomic predictions for drought tolerance in maize by joint modeling of additive and dominance effects in multi-environment trials. Heredity 121:24–37. https://doi.org/10.1038/s41437-018-0053-6

Dias KOG, Santos JPR, Krause MD, et al (2022) Leveraging probability concepts for cultivar recommendation in multi-environment trials. Theoretical and Applied Genetics 135:1385–1399. https://doi.org/10.1007/s00122-022-04041-y

Diepenbrock CH, Tang T, Jines M, et al (2022) Can we harness digital technologies and physiology to hasten genetic gain in us maize breeding? Plant Physiology 188(2):1141–1157. https://doi.org/10.1093/plphys/kiab527

Dunnington D (2023) Ggspatial: spatial data framework for ggplot2. URL https://CRAN.R-project.org/package=ggspatial, r package version 1.1.8

Eberhart SA, Russell WA (1966) Stability parameters for comparing varieties. Crop Science 6:36–40. https://doi.org/10.2135/cropsci1966.0011183X000600010011x

Ecmwf (2023) European centre for medium-range weather forecasts. URL https://www.ncei.noaa.gov/access/metadata/landing-page/bin/iso?id=gov.noaa.ncdc:C00765/

van Eeuwijk FA, Bustos-Korts DV, Malosetti M (2016) What should students in plant breeding know about the statistical aspects of genotype × environment interactions? Crop Science 56(5):2119–2140. https://doi.org/10.2135/cropsci2015.06.0375

Eosdis (2023) Nasa earth observing system data and information system. URL https://worldview.earthdata.nasa.gov

Fao (2014) World reference base for soil resources 2014. URL http://www.fao.org/3/i3794en/I3794en.pdf

Fick SE, Hijmans RJ (2017) Worldclim 2: new 1-km spatial resolution climate surfaces for global land areas. Int J Climatol 32:4302–4315. https://doi.org/10.1002/joc.5086

Finlay K, Wilkinson G (1963) The analysis of adaptation in a plant-breeding programme. Australian Journal of Agricultural Research 14:742. <https://doi.org/https://pdf.usaid.gov/pdfdocs/PNAAS139.pdf

Gauch JR HG, Zobel R (1997) Identifying mega-environments and targeting genotypes. Crop Science 37:311–326. https://doi.org/10.2135/cropsci1997.0011183X003700020002x

Ghcnd (2023) Global historical climatology network daily. URL https://www.ncei.noaa.gov/products/land-based-station/global-historical-climatology-network-daily/

Gilmour AR, Cullis B, Verbyla Ap (1997) Accounting for natural and extraneous variation in the analysis of field experiment. Journal of Agricultural, Biological and Environmental Statistics 2:269–293. https://doi.org/10.2307/1400446

Gogel B, Smith A, Cullis B (2018) Comparison of a one- and two-stage mixed model analysis of australia’s national variety trial southern region wheat data. Euphytica 214:44. https://doi.org/10.1007/s10681-018-2116-4

Guarino L, Jarvis A, Hijmans RJ, et al (2002) Geographic information systems (GIS) and the conservation and use of plant genetic resources. In: Managing plant genetic diversity. Proceedings of an international conference, Kuala Lumpur, Malaysia, 12–16 June 2000, CABI publishing Wallingford UK, pp 387–404

Guo Y, Xiang H, Li Z, et al (2021) Prediction of rice yield in East China based on climate and agronomic traits data using artificial neural networks and partial least squares regression. Agronomy 11(2):282. https://doi.org/10.3390/agronomy11020282

Heinemann AB, Costa-Neto G, Fritsche-Neto R, et al (2022) Enviromic prediction is useful to define the limits of climate adaptation: a case study of common bean in brazil. Field Crops Research 286:108628. https://doi.org/10.1016/j.fcr.2022.108628

Henderson CR (1949) Estimates of changes in herd environment. Journal of Dairy Science 61:294–300

Henderson CR (1950) Estimation of genetic parameters. Annals of Mathematical Statistics 21:309–310

Hernández MV, Ortiz-Monasterio I, Pérez-Rodríguez P, et al (2019) Modeling genotype × environment interaction using a factor analytic model of on-farm wheat trials in the yaqui valley of mexico. Agronomy Journal 111(6):2647–2657. https://doi.org/10.2134/agronj2018.06.0361

Hijmans R (2020) Raster: Geographic data analysis and modeling. R package version 3.6-3. URL https://CRAN.R-project.org/package=raster

Hijmans RJ, Barbosa M, Ghosh A, et al (2023) geodata: download geographic data. URL https://CRAN.R-project.org/package=geodata, r package version 0.5-8

Jarquin D, de Leon N, Romay C, et al (2021) Utility of climatic information via combining ability models to improve genomic prediction for yield within the genomes to fields maize project. Frontiers in Genetics 11:592769. https://doi.org/10.3389/fgene.2020.592769

Jarquíin D, Crossa J, Lacaze X, et al (2014) A reaction norm model for genomic selection using highdimensional genomic and environmental data. Theoretical and Applied Genetics 127(3):595–607. https://doi.org/10.1007/s00122-013-2243-1

Krause MD, Dias KOG, Singh AK, et al (2022) Using large soybean historical data to study genotype by environment variation and identify mega-environments with the integration of genetic and non-genetic factors. bioRxiv 4:487885. https://doi.org/10.1101/2022.04.11.487885

Lembrechts JJ, van den Hoogen J, Aalto J, et al (2022) Global maps of soil temperature. Global Change Biology 28(9):3110–3144. https://doi.org/10.1111/gcb.16060

Li X, Guo T, Mu Q, et al (2018) Genomic and environmental determinants and their interplay underlying phenotypic plasticity. Proceedings of the National Academy of Sciences 115(26):6679–6684. https://doi.org/10.1073/pnas.1718326115

Liland KH, Mevik BH, Wehrens R (2022) PLS: partial least squares and principal component regression. URL https://CRAN.R-project.org/package=pls, r package version 2.8-1

Lindgren F, Geladi P, Wold S (1993) The kernel algorithm for PLS. Journal of Chemometrics 7(1):45–59. https://doi.org/10.1002/cem.1180070104

Lynch M, Walsh B (1998) Genetics and analysis of quantitative traits, 1st edn. Sinauer Associates,

Sunderland Malosetti M, Ribaut JM, Eeuwijk FAV (2013) The statistical analysis of multi-environment data: modeling genotype-by-environment interaction and its genetic basis. Genetics Selection Evolution 4:44. https://doi.org/10.3389/fphys.2013.00044

Meuwissen THE, Hayes BJ, Goddard ME (2001) Prediction of total genetic value using genome-wide dense marker maps. Genetics 157:1819–1829. https://doi.org/11290733

Millet EJ, Kruijer W, Coupel-Ledru A, et al (2019) Genomic prediction of maize yield across european environmental conditions. Nature Genetics 51(6):952–956. https://doi.org/10.1038/s41588-019-0414-y

Montesinos-López OA, Montesinos-López A, Kismiantini ARoman-Gallardo, et al (2022a) Partial least squares enhances genomic prediction of new environments. Frontiers in Genetics 13:920689. https://doi.org/10.3389/fgene.2022.920689

Montesinos-López OA, Montesinos-López A, Sandoval DAB, et al (2022b) Multi-trait genome prediction of new environments with partial least squares. Frontiers in Genetics 13:966775. https://doi.org/10.3389/fgene.2022.966775

Monteverde E, Gutierrez L, Blanco P, et al (2019) Integrating molecular markers and environmental covariates to interpret genotype by environment interaction in rice (Oryza sativa L.) grown in subtropical areas. G3 Genes|Genomes|Genetics 9(5):1519–1531. https://doi.org/10.1534/g3.119.400064

Mrode RA (2014) Linear models for the prediction of animal breeding values, 3rd edn. CABI

NasaPower (2022) Prediction of worldwide energy resource. URL https://power.larc.nasa.gov/data-access-viewer

Ncei (2018) Climate forecast system reanalysis (CFSR), for 1979 to 2011. URL https://www.ncei.noaa.gov/access/metadata/landing-page/bin/iso?id=gov.noaa.ncdc:C00765/

Noaa (2023) Climate data online. URL https://www.ncei.noaa.gov/cdo-web

Nuvunga JJ, Silva CP, Oliveira LA, et al (2019) Bayesian factor analytic model: An approach in multiple environment trials. PLoS ONE 14(8):e0220290. https://doi.org/10.1371/journal.pone.0220290

Oliveira IC, Guilhen JHS, Ribeiro PCO, et al (2020) Genotype-by-environment interaction and yield stability analysis of biomass sorghum hybrids using factor analytic models and environmental covariates. Field Crops Research 257:107929. https://doi.org/10.1016/j.fcr.2020.107929

Ortiz R, Crossa J, Vargas M, et al (2007) Studying the effect of environmental variables on the genotype× environment interaction of tomato. Euphytica 153:119–134. https://doi.org/10.1007/s10681-006-9248-7

Ortiz R, Reslow F, Montesinos-López A, et al (2023) Partial least squares enhance multi-trait genomic prediction of potato cultivars in new environments. Scientific Reports 13(1):9947. https://doi.org/10.1038/s41598-023-37169-y

Patterson HD, Thompson R (1971) Recovery of inter-block information when block sizes are unequal. Biometrika 58:545–554. https://doi.org/10.2307/2334389

Pebesma E, Bivand R (2023) Spatial data science: with applications in R. URL https://r-spatial.org/book/

Piepho HP (1997) Analysis of a randomized block design with unequal subclass numbers. Agronomy Journal 89:718–723. https://doi.org/10.2134/agronj1997.00021962008900050002x

Piepho HP (2019) A coefficient of determination (r^2^) for generalized linear mixed models. Biometrical Journal 61(4):860–872. https://doi.org/10.1002/bimj.201800270

Piepho HP, Möhring J, Melchinger AE, et al (2008) Blup for phenotypic selection in plant breeding and variety testing. Euphytica 161:209–228. https://doi.org/10.1007/s10681-007-9449-8

Porker K, Coventry S, Fettell N, et al (2020) Using a novel PLS approach for envirotyping of barley phenology and adaptation. Field Crops Research 246:107697. https://doi.org/10.1016/j.fcr.2019.107697

R Core Team (2023) R: A Language and Environment for Statistical Computing. R Foundation for Statistical Computing, Vienna, Austria, URL https://www.R-project.org/

Ramburan S, Zhou M, Labuschagne M (2012) Integrating empirical and analytical approaches to investigate genotype × environment interactions in sugarcane. Crop Science 52(5):2153–2165. https://doi.org/10.2135/cropsci2012.02.0128

Resende RT, Piepho HP, Rosa GJM, et al (2021) Enviromics in breeding: applications and perspectives on envirotypic-assisted selection. Theoretical and Applied Genetics 134:95–121. https://doi.org/10.1007/s00122-020-03684-z

Rincent R, Malosetti M, Ababaei B, et al (2019) Using crop growth model stress covariates and AMMI decomposition to better predict genotype-by-environment interactions. Theoretical and Applied Genetics 132(12):3399–3411. https://doi.org/10.1007/s00122-019-03432-y

Rogers AR, Dunne JC, Romay C, et al (2021) The importance of dominance and genotype-by-environment interactions on grain yield variation in a large-scale public cooperative maize experiment. G3: Genes|Genomes|Genetics 11(2):jkaa050. https://doi.org/10.1093/g3journal/jkaa050

Sae-Lim P, Komen H, Kause A, et al (2014) Identifying environmental variables explaining genotype-by-environment interaction for body weight of rainbow trout (Onchorynchus mykiss): reaction norm and factor analytic models. Genetics Selection Evolution 46(16):1–11. https://doi.org/10.1186/1297-9686-46-16

Santos HG (2018) Sistema brasileiro de classificação de solos (in Portuguese), 5th edn. Embrapa, Brasília, DF, URL https://www.embrapa.br/en/busca-de-publicacoes/-/publicacao/1094003/sistema-brasileiro-de-classificacao-de-solos

Shelford VE (1911) Animal communities in temperate america as illustrated in the chicago region. Biological Bulletin 21:95–167. https://doi.org/10.5962/bhl.title.34437

Silva KJ, Teodoro PE, da Silva MJ, et al (2021) Identification of mega-environments for grain sorghum in Brazil using GGE biplot methodology. Agronomy Journal 113:1–12. https://doi.org/10.1002/agj2.20707

Smith A, Norman A, Kuchel H, et al (2021) Plant variety selection using interaction classes derived from factor analytic linear mixed models: models with independent variety effects. Frontiers in Plant Science 12:978248. https://doi.org/10.3389/fpls.2021.737462

Smith AB, Cullis BR (2018) Plant breeding selection tools built on factor analytic mixed models for multi-environment trial data. Euphytica 214:143. https://doi.org/10.1007/s10681-018-2220-5

Smith AB, Cullis B, Thompson R (2001) Analyzing variety by environment data using multiplicative mixed models and adjustments for spatial field trend. Biometrics 57:1138–1147. https://doi.org/10.1111/j.0006-341X.2001.01138.x

Smith AB, Ganesalingam A, Kuchel H, et al (2015) Factor analytic mixed models for the provision of grower information from national crop variety testing programs. Theoretical and Applied Genetics 128:55–72. https://doi.org/10.1007/s00122-014-2412-x

SoilGrids (2022) Soilgrids — global gridded soil information. URL https://www.isric.org/explore/soilgrids/

Sparks AH (2018) Nasapower: a nasa power global meteorology, surface solar energy and climatology data client for R. Journal of Open Source Software 3(30):1035. https://doi.org/10.21105/joss.01035

Stefanova KT, Buirchell B (2010) Multiplicative mixed models for genetic gain assessment in lupin breeding. Crop Science 50(3):880–891. https://doi.org/10.2135/cropsci2009.07.0402

Tolhurst DJ, Gaynor RC, Gardunia B, et al (2022) Genomic selection using random regressions on known and latent environmental covariates. Theoretical and Applied Genetics 135:3393–3415. https://doi.org/10.1007/s00122-022-04186-w

Van Eeuwijk FA, Elgersma A (1993) Incorporating environmental information in an analysis of genotype by environment interaction for seed yield in perennial ryegrass. Heredity 70(5):447–457. https://doi.org/10.1038/hdy.1993.66

Vargas M, Crossa J, Van Eeuwijk F, et al (2001) Interpreting treatment × environment interaction in agronomy trials. Agronomy Journal 93(4):949–960. https://doi.org/10.2134/agronj2001.934949x

Vargas M, van Eeuwijk FA, Crossa J, et al (2006) Mapping QTLs and QTL × environment interaction for CIMMYT maize drought stress program using factorial regression and partial least squares methods. Theoretical and Applied Genetics 112(6):1009–1023. https://doi.org/10.1007/s00122-005-0204-z

Wickham H (2016) Ggplot2: elegant graphics for data analysis. Springer. Springer. Cham, 2 editions

Wold HOA (1966) Estimation of principal components and related models by iterative least squares. Academic Press, New York

Wold S, Sjöström M, Eriksson L (2001) PLS-regression: a basic tool of chemometrics. Chemometrics and Intelligent Laboratory Systems 58:109–130. https://doi.org/10.1016/S0169-7439(01)00155-1

Wong J (2022) Pdist: partitioned distance function. URL https://CRAN.R-project.org/package=pdist, r package version 1.2.1

Wood J (1976) The use of environmental variables in the interpretation of genotype-environment interaction. Heredity 37(1):1–7. URL http://www.nature.com/articles/hdy197661

Xu Y (2016) Envirotyping for deciphering environmental impacts on crop plants. Theoretical and Applied Genetics 129:653–673. https://doi.org/10.1007/s00122-016-2691-5

Yan W, Hunt LA, Sheng Q, et al (2000) Cultivar evaluation and mega-environment investigation based on the GGE biplot. Crop Science 40:597–605. https://doi.org/10.2135/cropsci2000.403597x

Yan W, Kang MS, Ma B, et al (2007) GGE biplot vs. AMMI analysis of genotype-by-environment data. Crop Science 47:643–653. https://doi.org/10.2135/cropsci2006.06.0374

Yates F, Cochran WG (1938) The analysis of groups of experiments. Journal of Agricultural Science 28:556–580. https://doi.org/10.1017/S0021859600050978

