## Supplementary material for "GIS-FA: An approach to integrate thematic maps, factor-analytic and envirotyping for cultivar targeting": GIS-FA Supplementar 1

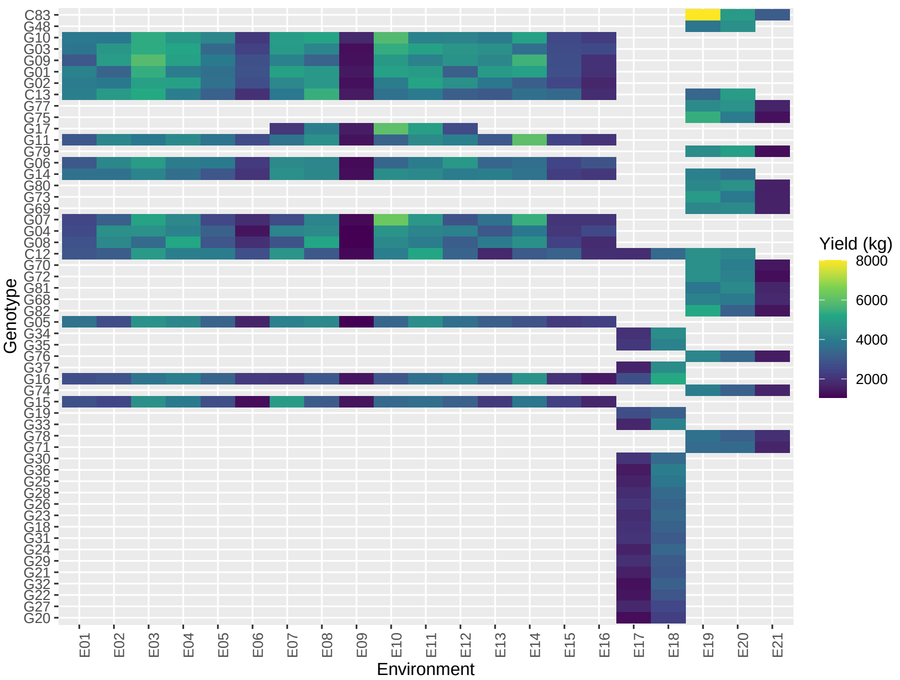


Supplementary Figure 1: Information about data balancing for rice and soybean cultivation. (a) corresponds to the Rice data evaluated during crop session 2009 and 2010

For the soybean dataset, we fitted thirteen statistical models to test if there are spatial trends within the plots (Supplementary Table 1). The standard model (M0) does not include any spatial adjustments. In the M1 model, the residual variance matrix was modeled by applying a first-order autoregressive structure ($AR1 \otimes AR1$) was applied in row and column directions. Many other structures were tested in the row direction, such as linear and spline. Because blocking was made in the column direction, we did not include the column effect in the model. Other models were also tested. Since the tested models have different fixed effects, the residual maximum likelihood cannot be compared using classical information criteria such as the Akaike Information Criterion (AIC) (Akaike, 1974). Therefore, we used the AIC corrected (Verbyla, 2019) ($AIC_c$), which can be calculated as follows:

$$AIC_c = -2logL + 2(p + q) \quad (1)$$

where $logL$ is the logarithm of the maximized value of the likelihood function for the fitted model, $p$ and $q$ are the number of fixed and random parameters in the model, respectively.

$^a lin(row)$ and $spl(row)$ are the effects of linear regression for the fixed effect of "row" and the spline component of "row" for the fixed effect, respec-

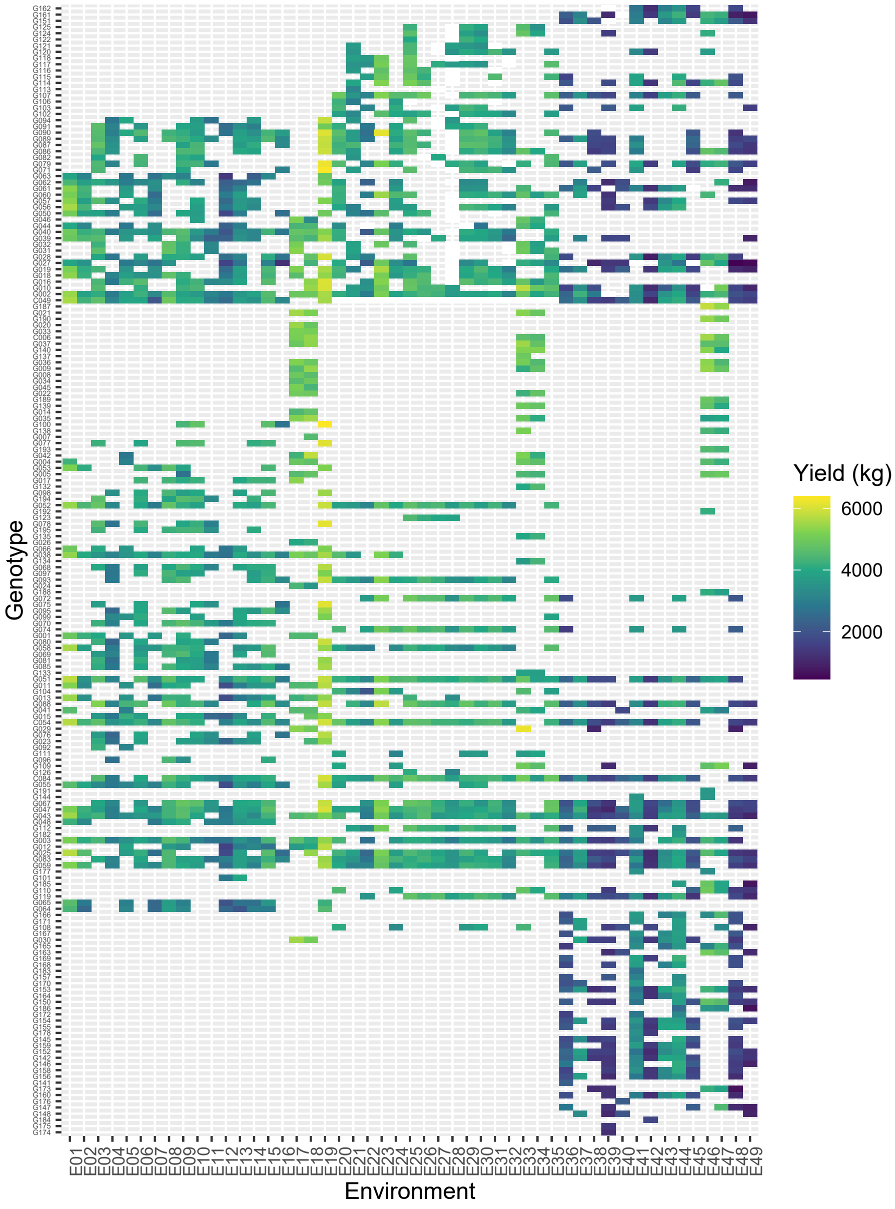


Supplementary Figure 1: Information about data balancing for rice and soybean cultivation. (b) Soybean data evaluated in 2019, 2020, and 2021. The colored cells indicate that the genotype was evaluated in that environment. The color corresponds to the productivity range. Cells without color represent the location where the genotype was not evaluated.

Supplementary Table 1: Evaluation locations of rice and soybean genotypes conducted in the state of Mato Grosso do Sul and in eight Brazilian states, respectively

| Rice data | | | | |
|---|---|---|---|---|
| **Location** | **State** | **2009/10** | | **2010/11** |
| Altamira | PA | E07 | | - |
| Anápolis | GO | E14 | | - |
| Boa Vista | RR | E01 | | - |
| Caceres | MT | E11 | | E20 |
| Colinas | MA | E04 | | - |
| Goianira | GO | - | | E17 |
| Marianópolis | TO | E05 | | - |
| Palmeiras de Goiás | GO | E15 | | - |
| Porto Nacional | TO | E06 | | - |
| Santa Carmem | MT | E10 | | - |
| Santo Antônio de Goiás | GO | E13 | | |
| São José dos Quatro Marcos | MT | - | | E21 |
| São Raimundo das Mangabeiras | MA | E02 | | E18 |
| Sinop | MT | E16 | | - |
| Sorriso | MT | E09 | | E19 |
| Teresina | PI | E03 | | - |
| Uruará | PA | E08 | | - |
| Vilhena | RO | E12 | | - |
| **Soybean data** | | | | |
| **Location** | **State** | **2019/20** | **2020/21** | **2021/22** |
| Anaurilândia | | E1, E2* | E20 | - |
| Antônio João | | E03 | E21 | E36 |
| Bela Vista | | - | - | E37 |
| Caarapó | | E04 | E22 | - |
| Campo Grande | | E05 | - | - |
| Itaporã | | E06 | E23 | E38 |
| Ivinhema | MS | E07 | E24 | E39, E40 |
| Maracaju | | E08, E09, E10, E11 | E25, E26, E27, E28 | E41 |
| Naviraí | | E12, E13 | E29, E30 | E42 |
| Rio Brilhante | | E14, E15, E16 | E31, E32 | E43, E44, E45 |
| São Gabriel do Oeste | | E17, E18 | E33, E34 | E46, E47 |
| Sidrolândia | | E19 | E35 | E48 |
| Terenos | | - | - | E49 |

*Trials conducted at the same location at different seasons.

Supplementary Table 2: Description of the 13 models adjusted for each productivity trial for the soybean dataset

| Model | Fixed effect[a] | Random effect[b] | Residual[c] |
|---|---|---|---|
| M0 | block | gen | $\mathbf{I}\sigma_e^2\mathbf{I}_r \otimes \mathbf{I}_c$ |
| M1 | block | gen | $\sigma_\xi^2[\sum_r(\rho_r) \otimes \sum_c(\rho_c)]$ |
| M2 | block | gen + nugget | $\sigma_\xi^2[\sum_r(\rho_r) \otimes \sum_c(\rho_c)]$ |
| M3 | block | gen + nugget + row | $\sigma_\xi^2[\sum_r(\rho_r) \otimes \sum_c(\rho_c)]$ |
| M4 | block + lin(row) | gen + nugget + row | $\sigma_\xi^2[\sum_r(\rho_r) \otimes \sum_c(\rho_c)]$ |
| M5 | block + spl(row) | gen + nugget + row | $\sigma_\xi^2[\sum_r(\rho_r) \otimes \sum_c(\rho_c)]$ |
| M6 | block + lin(row) | gen + nugget | $\sigma_\xi^2[\sum_r(\rho_r) \otimes \sum_c(\rho_c)]$ |
| M7 | block + spl(row) | gen + nugget | $\sigma_\xi^2[\sum_r(\rho_r) \otimes \sum_c(\rho_c)]$ |
| M8 | block | gen + row | $\sigma_\xi^2[\sum_r(\rho_r) \otimes \sum_c(\rho_c)]$ |
| M9 | block + lin(row) | gen + row | $\sigma_\xi^2[\sum_r(\rho_r) \otimes \sum_c(\rho_c)]$ |
| M10 | block + spl(row) | gen + row | $\sigma_\xi^2[\sum_r(\rho_r) \otimes \sum_c(\rho_c)]$ |
| M11 | block + lin(row) | | $\sigma_\xi^2[\sum_r(\rho_r) \otimes \sum_c(\rho_c)]$ |
| M12 | block + spl(row) | | $\sigma_\xi^2[\sum_r(\rho_r) \otimes \sum_c(\rho_c)]$ |

tively. [b]*row*: random effect of "row"; nugget $= \sigma_\eta^2(I_r \otimes I_c)$ is the variance-covariance structure of the independent part of the residual, assuming a first-order autoregressive correlation between rows and columns. [c]$AR1 \otimes AR1 = \sigma_\xi^2[\sum_r(\rho_r) \otimes \sum_c(\rho_c)]$ variance-covariance structure of the residual considers the first-order autoregressive correlation in the row-column direction, that is, the dependent portion of the residual; $\mathbf{I}$ is an identity matrix; $\sigma_e^2$ is the residual variance; $\sigma_\eta^2$ is the residual variance of the independent part; and $\sigma_\xi^2$ is the residual variance of the correlated part; *block* represents block repetition; *gen* represents the genotype. Although tested, the nugget effect $\eta$ was disregarded in the joint model due to convergence difficulties.

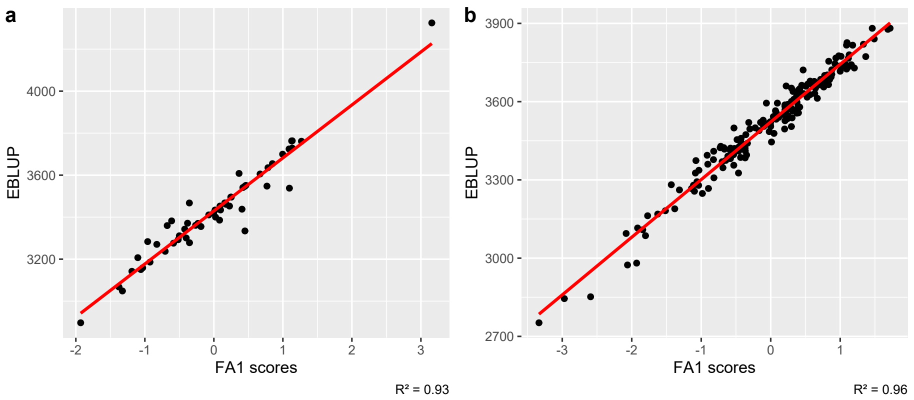


Supplementary Figure 2: Adjusted scores from the factor-analytic (FA) model displaying high correspondence plots between $FA_1$ scores and empirical Best Linear Unbiased Prediction (eBLUP) for (a) the Rice dataset and (b) the Soybean dataset

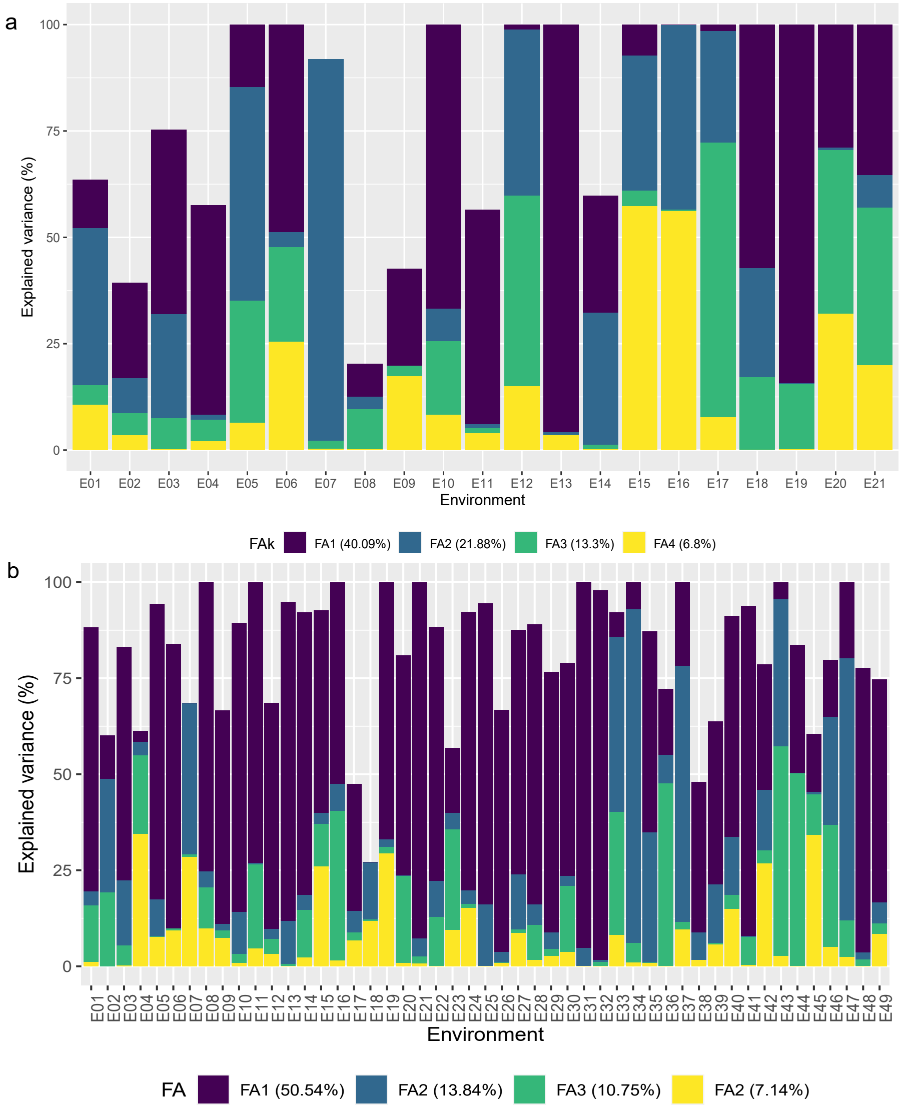


Supplementary Figure 3: Fitted factor analytic mixed models for the rice (a) and soybean (b) datasets and their total variance to four factors

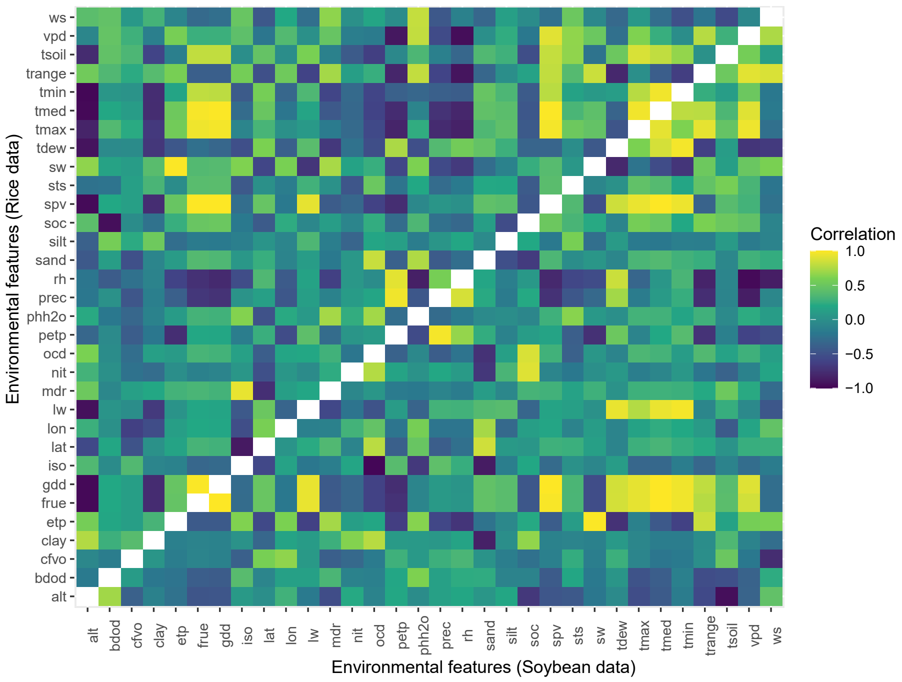


Supplementary Figure 4: Pearson correlation among 32 environmental features. On the left diagonal are all the features for the rice dataset, and on the right side are the soybean data

For the rice dataset, check C83 won all pure lines, so it was not plotted in the "who-wins-where" thematic map (Supplementary Figure 5).

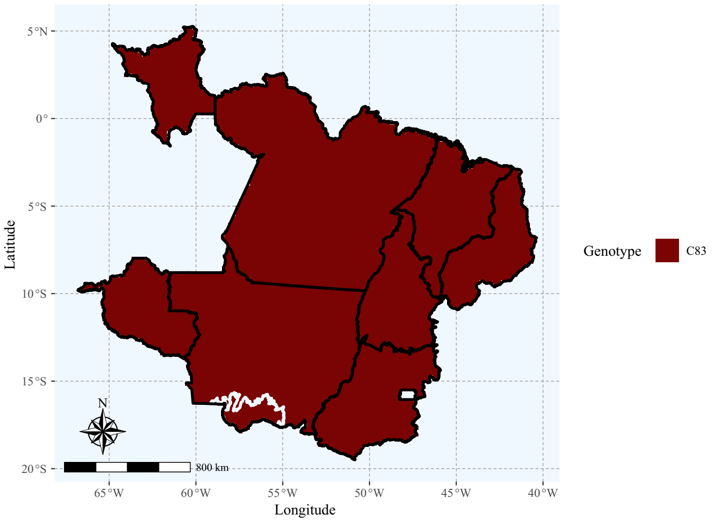


Supplementary Figure 5: Map wich-won-where with witness C83 from the rice dataset. This genotype was not compared with the others due to its superior productivity. The white line delimits the Pantanal biome
